# Social networks strongly predict the gut microbiota of wild mice

**DOI:** 10.1101/2020.09.24.311357

**Authors:** Aura Raulo, Bryony Allen, Tanya Troitsky, Arild Husby, Josh A Firth, Tim Coulson, Sarah CL Knowles

## Abstract

The mammalian gut teems with beneficial microbes, yet how hosts acquire these symbionts remains poorly understood. Research in primates suggests that microbes can be picked up via social contact, but the role of social interactions in non-group-living species remains unexplored. Here, we use a passive tracking system to collect high resolution spatiotemporal activity data from wild mice (*Apodemus sylvaticus*). Social network analysis revealed social association strength to be the strongest predictor of microbiota similarity among individuals, controlling for factors including spatial proximity and kinship, which had far smaller or nonsignificant effects. This social effect was limited to interactions involving males (male-male and male-female), implicating sex-dependent behaviours as driving processes. Social network position also predicted microbiota richness, with well-connected hub individuals having the most diverse microbiotas. Overall, these findings suggest social contact provides a key transmission pathway for gut symbionts even in relatively asocial mammals, that strongly shapes the adult gut microbiota. This work underlines the potential for individuals to pick up beneficial symbionts as well as pathogens from social interactions.

## Introduction

Symbiotic microbes are increasingly recognised as key modulators of host phenotypes. This is particularly true for the mammalian gut microbiota, whose metabolism is intimately entwined with that of the host. Among their many roles in host physiology, mammalian gut microbes modulate host energy metabolism (1,2), regulate fat accumulation and thermal homeostasis (3), and provide protection against pathogenic infection (4,5). They are also in constant dialogue with the host immune system, activating innate immune responses and tuning acquired immune responses to distinguish enemies from allies (6–8). As such, alterations to these microbial communities can have significant impacts on host health and have been associated with major metabolic and immune-related health conditions in humans (1,9,10)

Despite the well-established role of gut microbiota in host biology, we know surprisingly little about the forces that shape microbiota composition within and between individuals in nature. Community composition is notoriously variable among individuals, and is affected by a number of processes that can be viewed within a metacommunity framework (11): transmission processes (microbial dispersal) first determine which microbes colonize an individual host. Subsequently, aspects of the nutritional and immunological environment inside the host (e.g. host diet, genetics), as well as ecological interactions with resident microbes, selectively filter colonising microbes that can persist and thrive. In mammals, the microbiota is initially established through maternal transmission at birth (12), with community composition then further shaped by transmission from family members and the broader environment (13–15) as well as selective processes within the host (16).

A key question is to what extent ongoing transmission throughout life shapes the microbiota. Accumulating evidence suggests the gut microbiota is affected by a host’s environment, such as diet (17,18) and contact with soil (15,19,20). The microbiota can also be shaped by a host’s social environment, since a special form of microbial transmission can occur through social contact. Intimate social contact, such as the many forms of prosocial touch common in mammals (e.g. grooming, licking, huddling), may function as an important transmission route for microbes. This is particularly true for microbes not easily transmitted via the environment, including strict anaerobes and non-spore-forming bacteria (21). Moreover, if less transmissible microbes are more likely to positively impact host fitness (22), social interactions could constitute a key pathway (alongside vertical transmission) by which symbionts of high functional significance are transmitted in mammals. Laboratory rodent studies have repeatedly shown that cohousing drives convergence in microbiota composition (23–25), indicating that social interaction and close proximity facilitate microbial transmission.

In highly social group-living mammals, the host social environment seems to have important effects on the gut microbiota. Social group membership has been shown to predict gut microbiota composition in several species of primates (26–31) and other group-living mammals (32–34). Social group effects also occur in humans, as unrelated individuals living in the same household were found to have a more similar microbiota than relatives living in different households (35). However, the mechanisms underlying these observations remain unclear, and may include not only direct social transmission but also shared environmental exposures like diet. In some cases, social group effects on the microbiota have been found while controlling for kinship or shared diet, supporting the idea that social transmission homogenises the gut microbiota. For example, sifakas (*Propithecus verrauxii*) were found to have a social group-specific gut microbiota composition that was not explained by dietary or habitat overlap, nor genetic relatedness among group-members (28). Further support comes from individuals observed to switch social groups, for example immigrant male baboons (36), whose microbiota composition converges on that of their new social group.

Some evidence also suggests social interactions affect microbiota similarity at a dyadic level, within groups or populations. Several primate studies have shown the intensity of social interaction between group members to predict similarity in their microbiota (26–28,30). Baboons that groomed each other more were found to share more gut microbes, and these shared bacteria were enriched in anaerobic and non-spore forming taxa (26). Similar patterns were found in humans, with couples who reported having a “physically close relationship” sharing more gut microbes than less close couples or friends (37). However, socially interacting primates often experience strong overlap in their environments, and thus it remains difficult to distinguish social transmission from shared environmental exposures (21). Species that are not group-living (sensu Wilson, 38) arguably provide more powerful systems in which to clearly distinguish effects of social interaction from confounding shared environmental exposures, as social interactions are more limited in time and space. However, the role of social transmission in shaping the microbiota in such species has yet to be explored.

Here, we use wild mice as a model system (wood mice, *Apodemus sylvaticus*) to assess how social interactions shape gut microbiota similarity among sympatric mice, in comparison to effects of host kinship, spatial proximity, and other factors. These mice are not group-living, but can be considered a semi-social species, with the propensity to co-nest in underground burrows varying seasonally and between individuals (39,40). Individuals have stable, partially overlapping home ranges, and yet vary in their level of social contact, making them a particularly suitable species in which to study social transmission. Using a tracking system based on passive radio-frequency identification (RFID) tags, we intensively followed a population of mice for one year and used social network analyses to test two specific hypotheses about social transmission of microbiota. First, we test the prediction that if social interactions drive microbial transmission, dyadic microbiota similarity will be positively predicted by proximity in the social network, independent of other potential confounders. Second, individuals that are more connected in the social network are predicted to have higher microbiota diversity, as they are exposed to more extensive social transmission.

## Materials and Methods

### Field data collection

Data were collected over a one-year period (Nov 2014-Dec 2015) from a wild population of wood mice (*Apodemus sylvaticus*) in a 2.43ha mixed woodland plot (Nash’s Copse) at Imperial College’s Silwood Park campus, UK (Figure S1A). Live traps were set for one night every 2-4 weeks in an alternating checkerboard design, to ensure even coverage. At first capture, all mice were injected subcutaneously with a passive integrated transponder tag (PIT-tag) for permanent identification. At each trapping, demographic data on captured animals was recorded and samples for gut microbiota analysis and mouse genotyping collected (see Supplementary Material).

Data on rodent space use and social associations was collected in parallel to trapping using a set of 9 custom-built PIT-tag loggers (described in 41 and Supplementary Material; Figure S2), distributed across the trapping grid. Loggers consisted of a box with entrance tubes, that recorded the time-stamped presence of any rodent that entered. Loggers were rotated systematically around the plot throughout the study period, using a sampling design that ensured even spatial coverage, with each 100m^2^ grid cell covered on average 5.49 (SD 1.61) times (Figure S1B). Of the 93 mice tagged during study period, 89% (n=83) were detected by the loggers.

### Kinship analysis

To derive estimates of host genetic relatedness, ear tissue samples were used to genotype all captured mice at eleven microsatellite loci (Table S1, Table S2; detailed in 39) and build a pedigree in COLONY 2.0.6.5 (42). Full details of genotyping methods and pedigree reconstruction are provided in Supplementary Material. After sample failures, genetic relatedness could be inferred for 70 of the 83 monitored mice.

### Constructing social networks

All analyses were conducted in R version 3.6.1 (R-Core-Team 2019). To capture patterns of spatiotemporal coincidence among wood mice, social networks were constructed from logger data using the package *asnipe* (43) and plotted using *igraph* (44). Individual mice were nodes, and edges described the number of times two individuals were observed at the same logger with the same night (12h period, 6pm to 6am). To measure association strength, we used an adjusted version of the Simple Ratio Index (SRI), that accounted for variable overlap in individual lifespans (i.e. time between first and last logger observation) (45), hereafter “Adjusted SRI”. Adjusted SRI is defined as follows for two individuals, A and B:

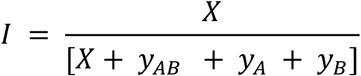

where *X* is the number of instances (night-location combinations) in which A and B were observed associated (observed within a specific time window of each other), *y*_*AB*_ is the number of instances in which *A* and *B* were both observed, but not associated, *y*_*A*_ and *y*_*B*_ are the number of instances in which both were known to be alive but only A or B were observed respectively. By taking lifespan overlap into account we could incorporate data from all 83 individuals across the entire year into one static social network.

To examine how the definition of social association might affect social network-microbiota relationships, we constructed a series of networks using increasingly intimate definitions of social association, by applying a sliding time window of variable length to define social association, from 12 hours (as above) down to a 2 minute period (12h, 4h, 1h, 30min, 10min, 2min). We also calculated a parallel set of networks with binary social association indices (BI), where ‘1’ indicated the dyad were observed associated at least once, and ‘0’ indicating they were not.

### Gut microbiota characterisation

The gut microbiota was successfully characterised from 239 faecal samples belonging to 75 individual wood mice (covering 90% of the monitored mice, mean=3.2 samples/mouse, range=1-9). Full details of library preparation, sequencing and bioinformatics are given in Supplementary Material. Briefly, microbiota profiling involved amplicon sequencing of the 16S rRNA gene (V4-region). Sequence data were processed through the DADA2 pipeline v1.6.0 (46), to infer amplicon sequence variants (ASVs) and taxonomy assigned using the GreenGenes Database (Consortium 13.8). Using package *phyloseq* (47), ASV-counts were normalized to proportional abundance within each sample (48) and singleton ASVs as well as those belonging to non-gut microbial taxa (Cyanobacteria, Mitochondria) were removed. Lastly, we used package iNEXT (49) to estimate asymptotic richness and Shannon diversity for each sample.

### Statistical analyses

To describe compositional microbiota variation, package *vegan* (50) was used to calculate Jaccard distances and Bray-Curtis dissimilarities among samples (Figure S7). We used the Jaccard Index (1-Jaccard distance, the proportion of shared ASVs between sample pairs) as our primary measure of microbiota similarity, as we considered this metric most relevant for investigating microbial transmission among hosts. However, we repeated key analyses using Bray-Curtis dissimilarity, an abundance-weighted metric less sensitive to potential sequencing artefacts.

#### General predictors of gut microbiota composition

We performed permutational analysis of variance (PERMANOVA) in *vegan* to 1) test the repeatability of gut microbiota composition among individuals sampled multiple times, 2) identify non-social effects on the microbiota that should be controlled for in subsequent analyses and 3) estimate how much individual variation was independent of these covariates. We tested effects of time (factor month), host age (juvenile/adult), sex, plot region, habitat type, and individual identity on Jaccard distance. Plot region and habitat type for each individual were defined from logger data, as the most common logger territory (no.1-9) and habitat type (rhododendron, open woodland/bluebell, bamboo or mixed; Figure S1) they were detected in.

#### Associations between social association strength and microbiota similarity

To test whether dyadic microbiota similarity was predicted by social association strength, we performed Bayesian regression models in package *brms* (51). These models are well-suited for this as they permit random effect structures able to account for the types of dependence inherent to dyadic data, and repeat sampling of individuals (52). We constructed *brms* models that included all dyadic sample comparisons except within-individual comparisons. Microbiota similarity (Jaccard Index) was used as the response, with social association strength (adjusted SRI, or BI index) as the main predictor. As the Jaccard Index is a proportion, a logit link function was used. To control for potential confounding variables as far as possible, we fitted several dyadic covariates: spatial distance between hosts, sampling interval (time in days between samples taken), kinship, sex and age similarity (0/1 for different/same). Spatial distance was calculated as the distance between individuals’ mean spatial coordinates from logger records (minimum 34 logger records per mouse). All covariates either naturally ranged from 0 to 1 or were scaled to do so, to make model estimates for all terms comparable. To control for non-independence in the dataset arising from a dyadic response variable and repeat samples per mouse, both the model intercept and slope (social association strength effect) were allowed to vary as defined by two random effects: i) a multi-membership random effect capturing the individuals in each dyad (Individual A + Individual B) and ii) a multi-membership random effect capturing the samples in each dyad (Sample A + Sample B).

To test for sex-dependence in the effect of social association (e.g. arising from specific sexual behaviours) on microbiota, the main model (12h edge definition) was also run including dyad sex category (male-male, male-female or female-female) and its interaction with social association strength. In this model, only a multi-membership random intercept was fitted (not a random slope) to help ensure there was enough power to estimate the interaction effect. Finally, to check our results were robust to the chosen statistical approach, we confirmed key results with two alternative statistical modelling frameworks: 1) *MCMCglmm*, an alternative R package for Bayesian regression (53) and 2) a matrix permutation-based method common in social network analyses, Multiple Regression Quadratic Assignment procedure (MRQAP; 54), with a data subset including one randomly selected sample per individual (Supplementary Material).

#### Social network position and microbiota diversity

We hypothesized that an individual’s social network position might affect gut microbiota (alpha) diversity. Depending on the transmission ecology, different types of network position might best predict diversity. To explore this, we calculated six different metrics of network position, that capture different aspects of social connectedness (Figure 1). If the sheer amount of social interaction or number of social partners can diversify the microbiota, we expect diversity to be predicted by measures of general network centrality (Figure 1). Alternatively, if diversity is driven by the distinctness of transmission sources, and if this is reflected in their social distance, we expect diversity to be predicted by measures of bridge-type centrality (Figure 1). To test the relationship between each centrality measure and gut microbiota diversity, we used Bayesian regression models in *MCMCglmm* with either asymptotic ASV richness or asymptotic Shannon diversity as the response. We first explored how several covariates predicted diversity: host age, sex, sampling month (as a factor), plot region, habitat, read count, and PCR plate (4-level factor), and simplified models to include only covariates with p<0.1. We then added into the model one of our six measures of social centrality (Figure 1), derived from either the 12h or 2min network. Individual identity and PCR plate were fitted as a random factors. A node permutation test was used to verify that significant effects were not driven by network structure. Here the observed posterior mean estimates for network position were compared with those derived from 1000 models in which network positions were randomised across individuals.

**Figure 1:**
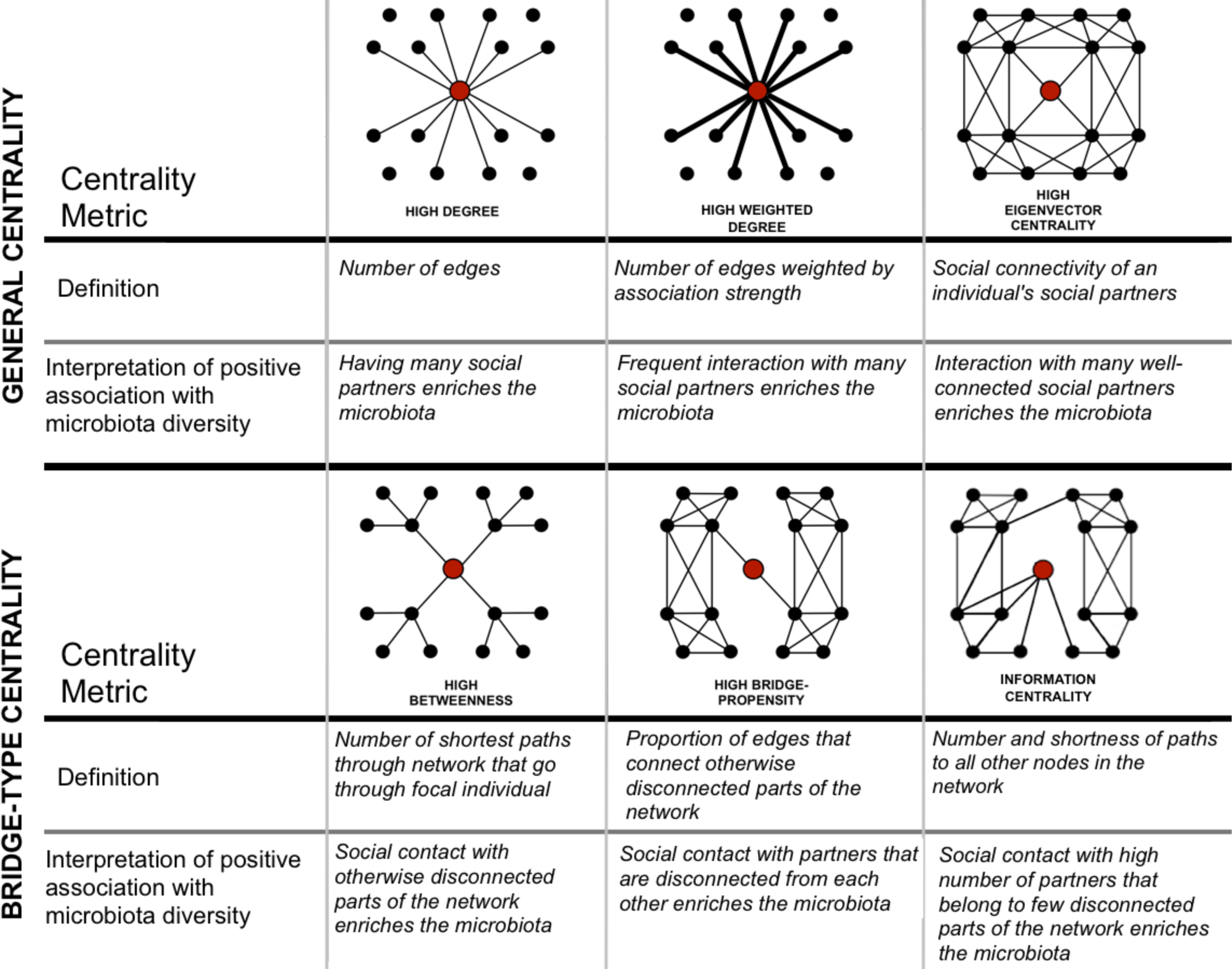
Six measures of network centrality and interpretation of a positive relationship with microbiota diversity. Images depict focal individuals (red circles) whose social interactions (lines) with other individuals (black circles) give them a high value of each centrality metric.

#### Identifying which bacterial taxa associate with social interaction

To identify candidate socially transmitted bacterial taxa, we tested how each bacterial family affected the strength of correlation between social association strength and microbiota similarity. We recalculated the Jaccard Index excluding each bacterial family in turn, then compared (both 12h and 2min) social network effect sizes and credible intervals from MCMCglmm models using these indices (full model details in Supplementary Material).

## Results

### Factors predicting gut microbiota composition

In a marginal PERMANOVA on data from repeat-sampled mice, individual identity explained 33% compositional variation in the microbiota, while temporal fluctuations (month) explained 6%, with similar results for both Jaccard Index and Bray-Curtis dissimilarity (Table S3). When other individual-level attributes were included (age, sex, plot region and habitat type), 27% variation in microbiota composition remained attributable to individual identity (Table S4), indicating the microbiota showed consistent individual variation that was not explained by measured host factors. No other variables predicted microbiota composition, except for a weak effect of habitat type (marginal PERMANOVA on data with one sample per individual, Table S5). Among the subset of hosts (70 of 75) with kinship information, kinship and microbiota similarity (Jaccard Index) were unrelated (Mantel test: r=0.001, p=0.520).

### Wood mouse social structure

The wood mouse social network showed marked variation in edge weights (social association strength) but no clear clustering, and global network density declined as increasingly intimate edge definitions were used (Figure 2A-D). Consequently, the correlation among social networks with different edge definitions decayed as the difference in time windows increased (Table S6). As expected, social association strength was to an extent predicted by spatial proximity in all networks (MRQAP p<0.001, Table S7), though this spatial effect weakened as more intimate edge definitions were used (Figure S9). Even in the least intimate (12h) social network, mice clearly did not solely associate with their nearest neighbours, as distances to the closest social partner (mean 25.6m; sd=15.3m) were on average over three times greater than those to the nearest neighbour (mean=8.4m; sd=5.5m). Some strong social associations were observed between individuals whose mean spatial locations were over 60 meters apart (Figure 2E-H). As such, the social structure of this population was only partially determined by spatial location, and this spatial influence on social contact was weakest in the 2min network.

**Figure 2:**
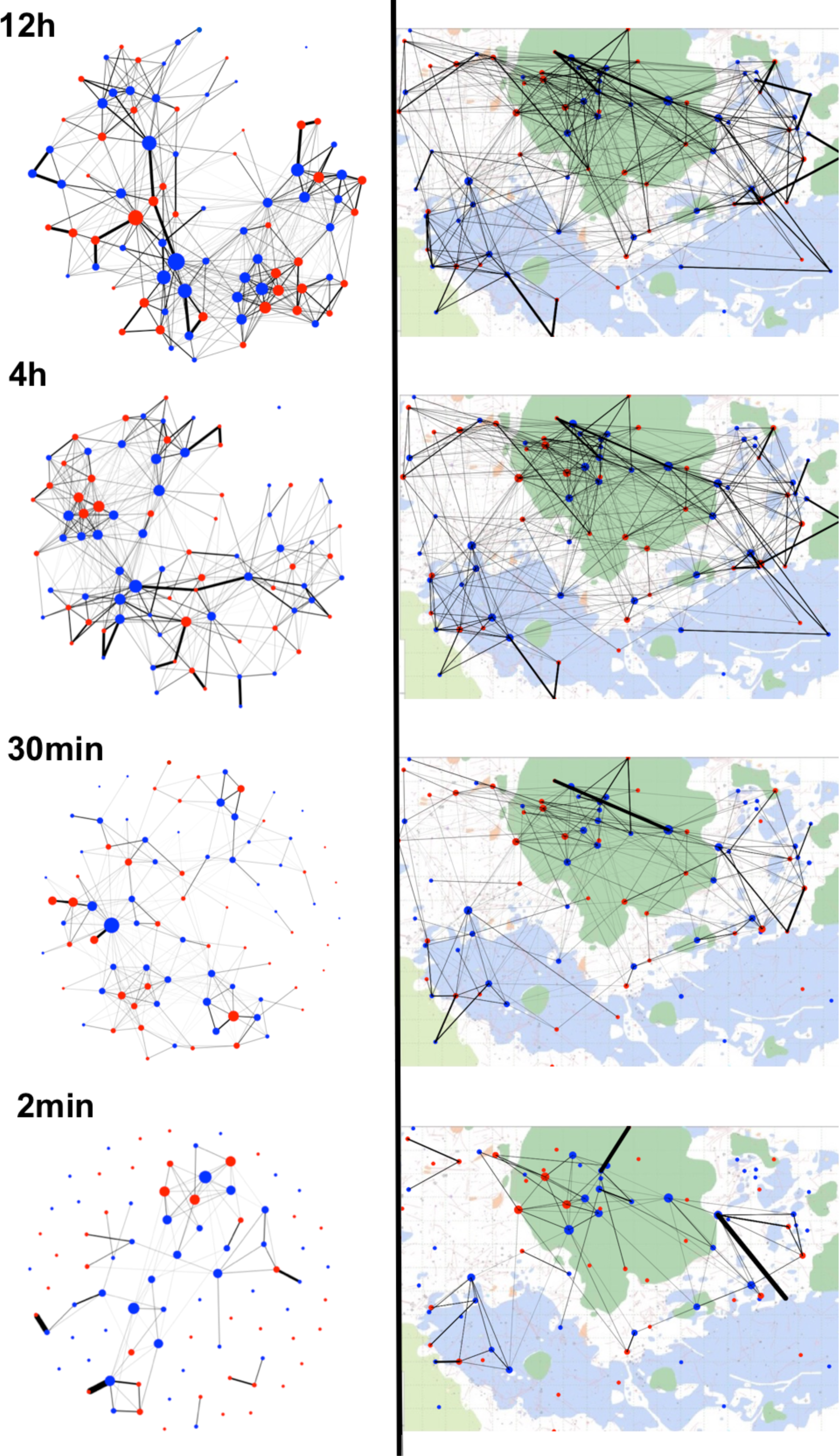
Wild wood mouse social networks plotted in either (A) social space or (B) geographical space. In A) networks are plotted using a standard weighted spring layout that minimises the sum of edge lengths and overlap across the network (*igraph*, (43)), and in B) mice are positioned at their mean spatial coordinates recorded from logger data, superimposed on a habitat map of the study area. Background colours reflect habitat types (dark green=rhododendron, light green=bamboo, blue= bluebell, white= open woodland). Red and blue circles represent female and male mice respectively, and line thickness is proportional to social association strength.

### Social association strength predicts microbiota similarity

Among pairs of individuals, the strength of social association strongly and positively predicted similarity in gut microbiota composition (in 12h network: Posterior mean 0.78, CI=0.34 to 1.24; Figure 3). Specifically, the proportion of ASVs shared within dyads (Jaccard Index) was positively predicted by their social association strength in all networks, even when controlling for effects of sex, age, kinship, sampling interval, and spatial distance (Table S8). Other variables also predicted microbiota similarity, including the spatial distance between hosts (Posterior Mean -0.08, CI=-0.12 to -0.04) and the time interval between which they were sampled (Posterior Mean -0.46, CI= -0.48 to -0.43), but the size of these effects was consistently smaller than that of social association strength (Figure 3, Table S8). Similar results were obtained from models using alternative statistical frameworks, or using Bray-Curtis dissimilarity instead of the Jaccard Index (Supplementary Material). Even binary social networks predicted microbiota similarity (Table S11), albeit less strongly than association strength.

**Figure 3:**
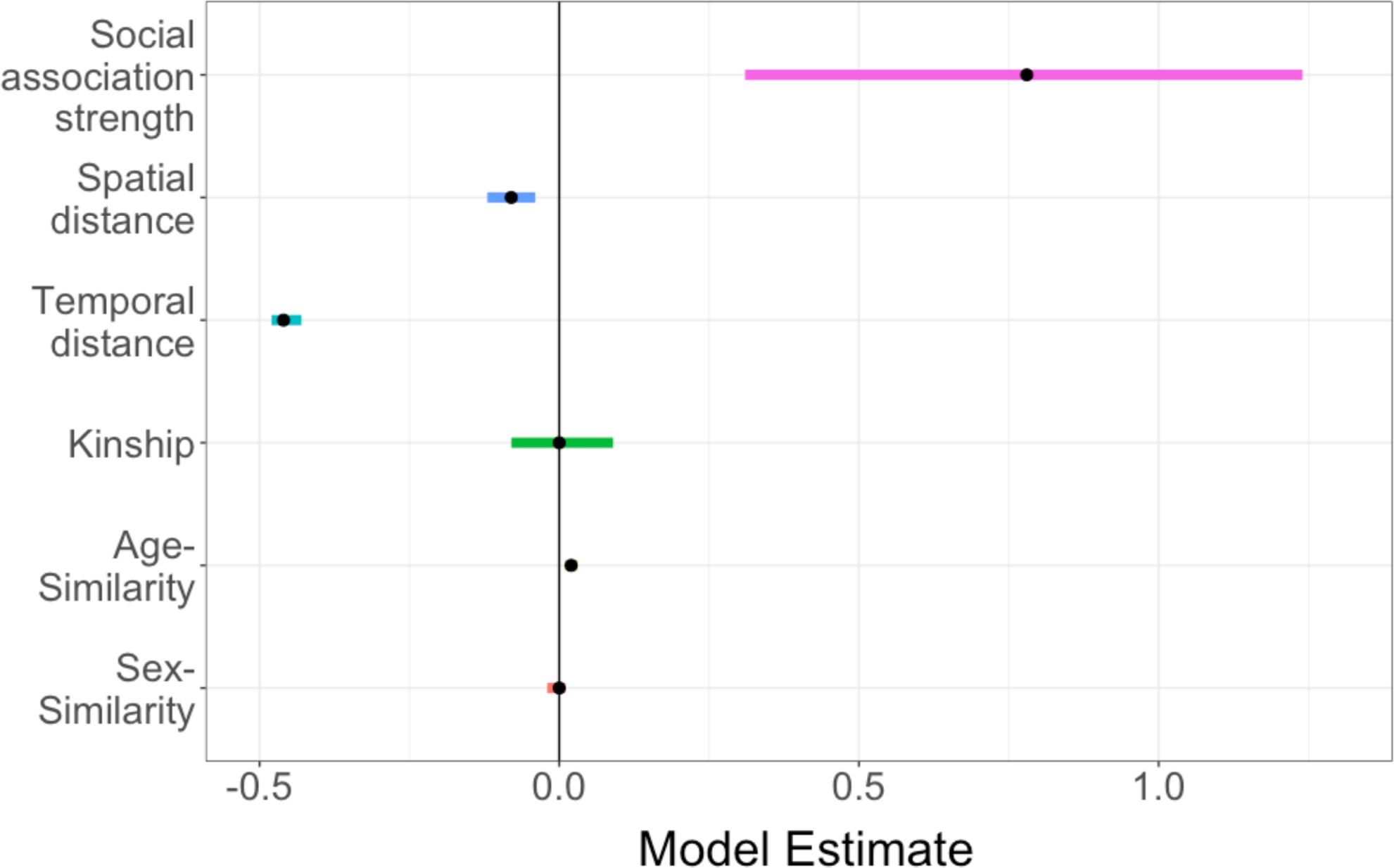
Social association strength predicts gut microbiota similarity more strongly than spatial distance, kinship and other effects. Effect size estimates (points) and their 95% credible intervals (coloured lines) are plotted from Bayesian regression (*brms*) models with pairwise microbiota similarity among hosts (Jaccard Index) as the response. Where confidence intervals do not overlap zero, a variable significantly predicts microbiota similarity. Social association strength in the 12h network has a strong positive effect on microbiota similarity, that is larger than that of other variables.

The relationship between social association strength and microbiota similarity became stronger as networks with increasingly intimate edge definitions were analysed (Figure 4A), while spatial and temporal effects remained comparable across networks (Table S8). As such, the effect of social association increased from 1.7 times as large as the next strongest (temporal) effect in the 12h network, to over 13 times as strong in the most intimate (2min) network. Since more intimate networks also had fewer edges (i.e. lower density, Figure 2), we also tested whether variation in network density alone might drive this trend. To do this, we ran a set of null models (described fully in Supplementary Material) in which the least intimate (12h) network was thinned to have the same number of edges as seen in each real network. In contrast to the real networks, social network effect sizes remained relatively constant in null models using artificially thinned networks (Figure 4B).

**Figure 4:**
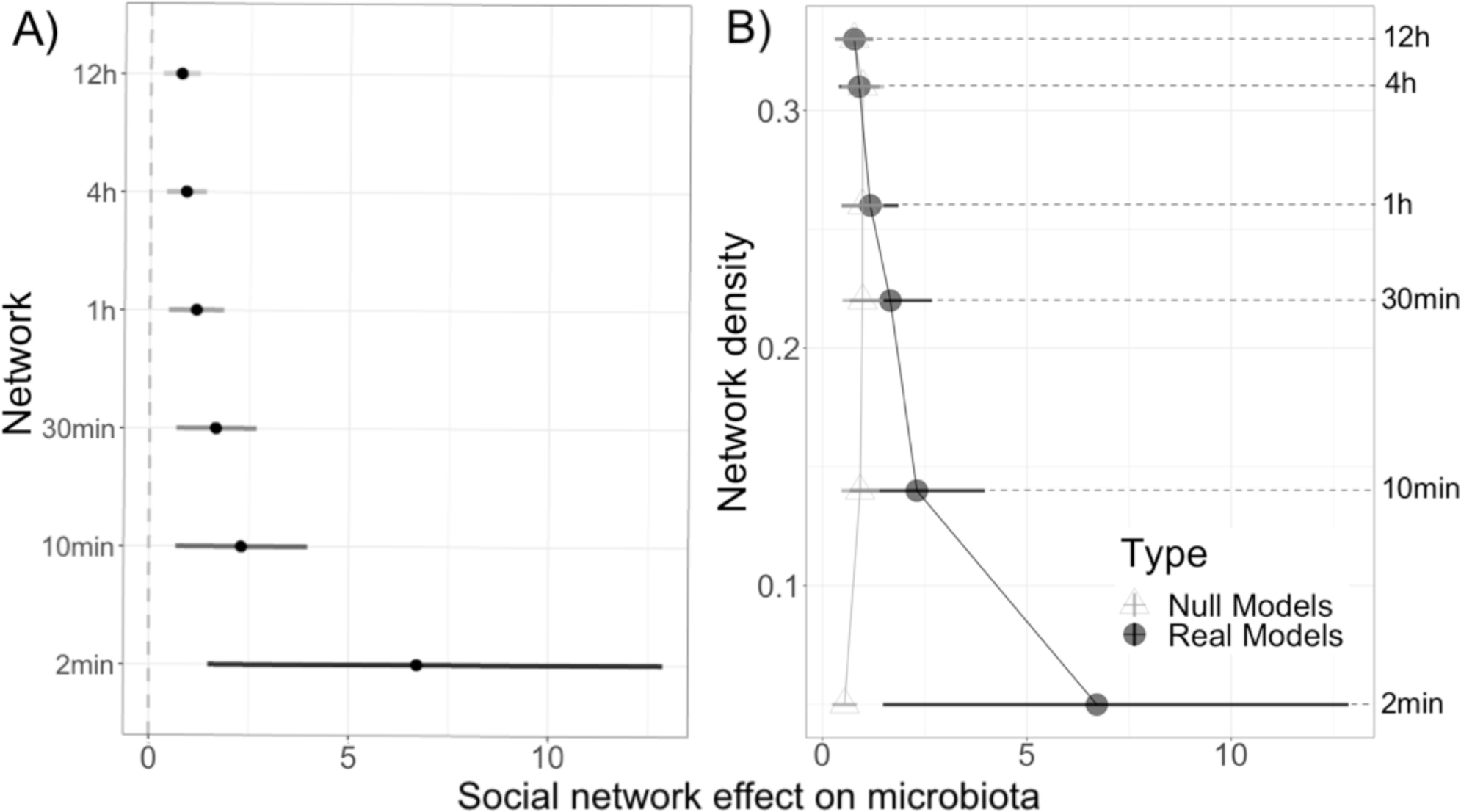
Social association strength predicts microbiota similarity more strongly in networks that use a more intimate edge definition. A) The effect of social association strength on microbiota similarity (Jaccard Index) is stronger in networks with more intimate edge definitions. Social network effect sizes (estimated slope of the relationship between social association strength and microbiota similarity, the Jaccard Index) and their 95% credible intervals are plotted from Bayesian regression (*brms*) models that included the same covariates shown in Figure 3. B) Differences in effect size across networks are not due to variation in network density, as effect size did not change in null models where the 12h network was artificially thinned by removal of the weakest edges to have the same density as each real network of differing edge definition.

### Sex-dependent effects of social association on microbiota similarity

We further found that the effect of social association strength on microbiota similarity depended on the sex of interacting individuals. In a model including an interaction between social association strength and dyadic sex combination, social association strength predicted microbiota similarity strongly in male-male pairs (posterior mean 0.28, CI= 0.14 to 0.54; Table S10) and male-female pairs (posterior mean 0.29, CI= 0.04 to 0.55) but not significantly in female-female pairs (posterior mean 0.1, CI -0.14 to 0.34; Figure 5, Table S10).

**Figure 5:**
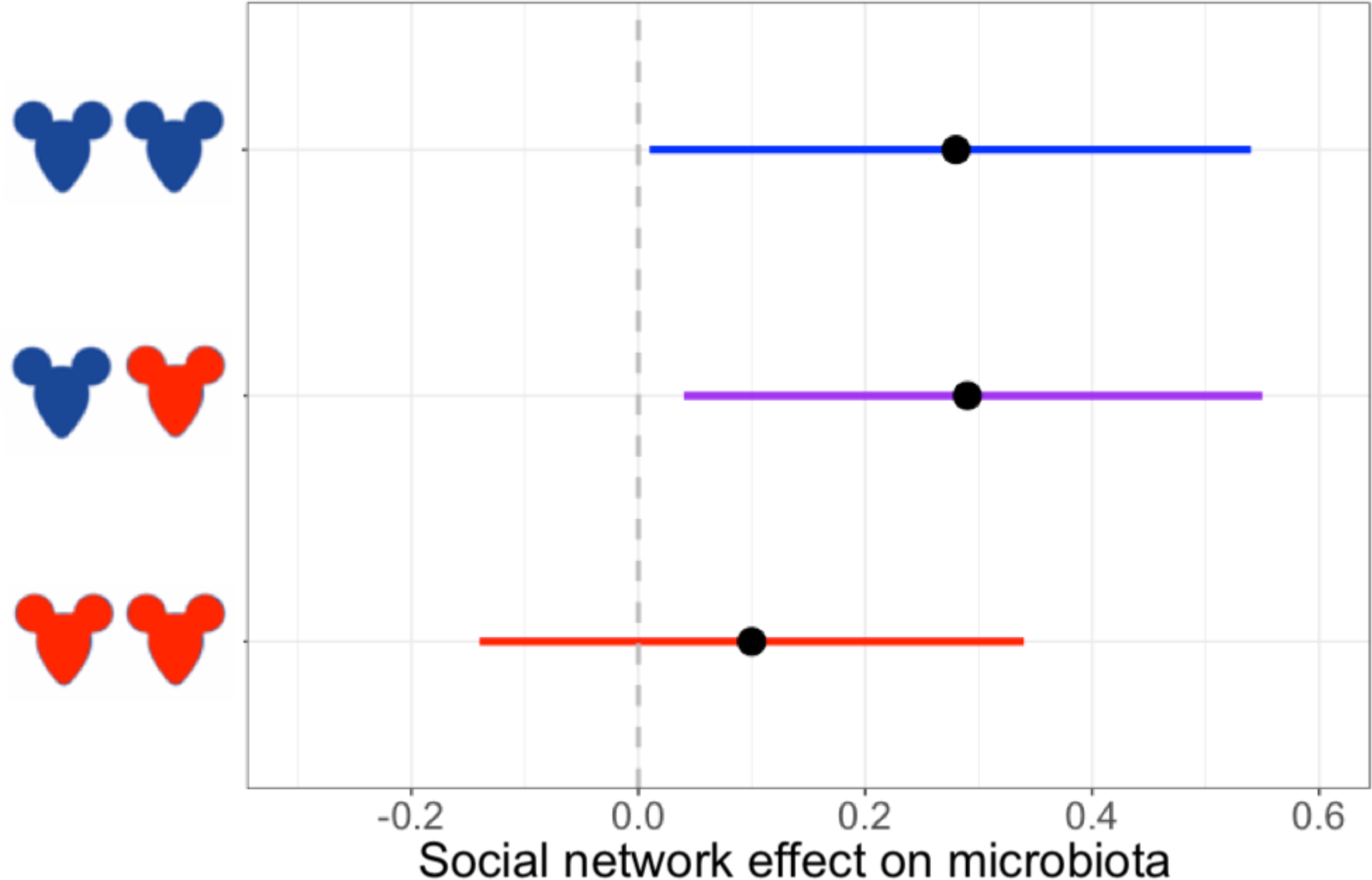
Social association strength predicts microbiota similarity only among dyads involving males. Estimated social network effects on the microbiota (slope of the relationship between social association strength and Jaccard Index) and 95% credible intervals are plotted from a Bayesian regression (*brms*) model predicting microbiome with the 12h social network that included an interaction term between social association strength and dyad sex-category (male-male, male-female or female-female). Females are depicted in red and males in blue respectively. Social association strength has a significant positive association with microbiota similarity in dyads involving males, but not in female-only dyads.

### Social network position and microbiota diversity

Both microbiota diversity metrics (richness and Shannon diversity) were predicted by plot region, habitat type, and month (Table S13). Both diversity estimates were also associated with PCR plate, and richness was also predicted by read count. Four measures of network position positively predicted gut microbiota richness: degree and information centrality predicted richness in both 12h and 2min networks, and betweenness and bridge propensity additionally predicted richness in the 2min network (Table 1). No measures of network position predicted Shannon diversity when controlling for covariates (Table S12).

**Table 1:**
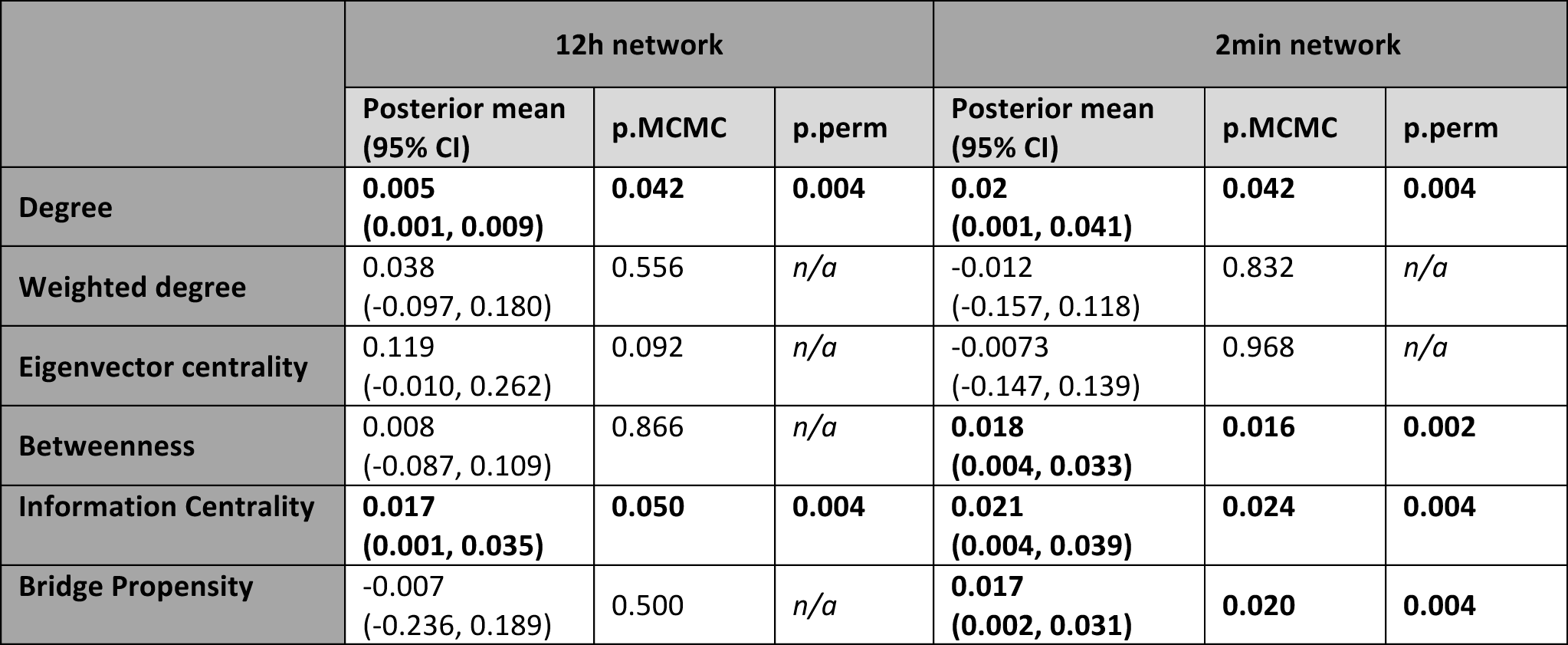
Social network centrality metrics predict individual gut microbiota richness. Posterior means and 95% credible intervals are shown from MCMCglmm models including the covariates shown in Table S13 and a single centrality metric. Significant effects are shown in bold. Significance was inferred from two p-values: If the Bayesian model p-value calculated from posterior distribution (p.MCMC) <0.05, the result was further tested by calculating a permutational p-value (p.perm). p.perm represents the probability of generating the observed posterior mean given the data, based on 1000 node-based permutations in which the centrality values of nodes are randomly shuffled before running the model.

### Identifying bacterial taxa that drive social network effects

The social network effect we identified did not depend entirely on any single bacterial family, since it remained statistically significant in all models where a single bacterial family was excluded (Figure 6). For some of the more diverse bacterial families, effect size did shift when they were excluded, but not in a way that directly related to their diversity. Excluding the family S24-7 made the social network effect weaker and almost non-significant when using the most intimate (2min) edge definition (taking the p.MCMC-value from p<0.001 to p=0.012), a pattern that was similar but weaker in the 12min network. Conversely, excluding Lachnospiraceae, the most diverse family, if anything slightly strengthened the social network effect in both networks (Figure 6). Excluding Lactobacillaceae also weakened the social network effect size somewhat, but only when using the less intimate (12h) edge definition.

**Figure 6:**
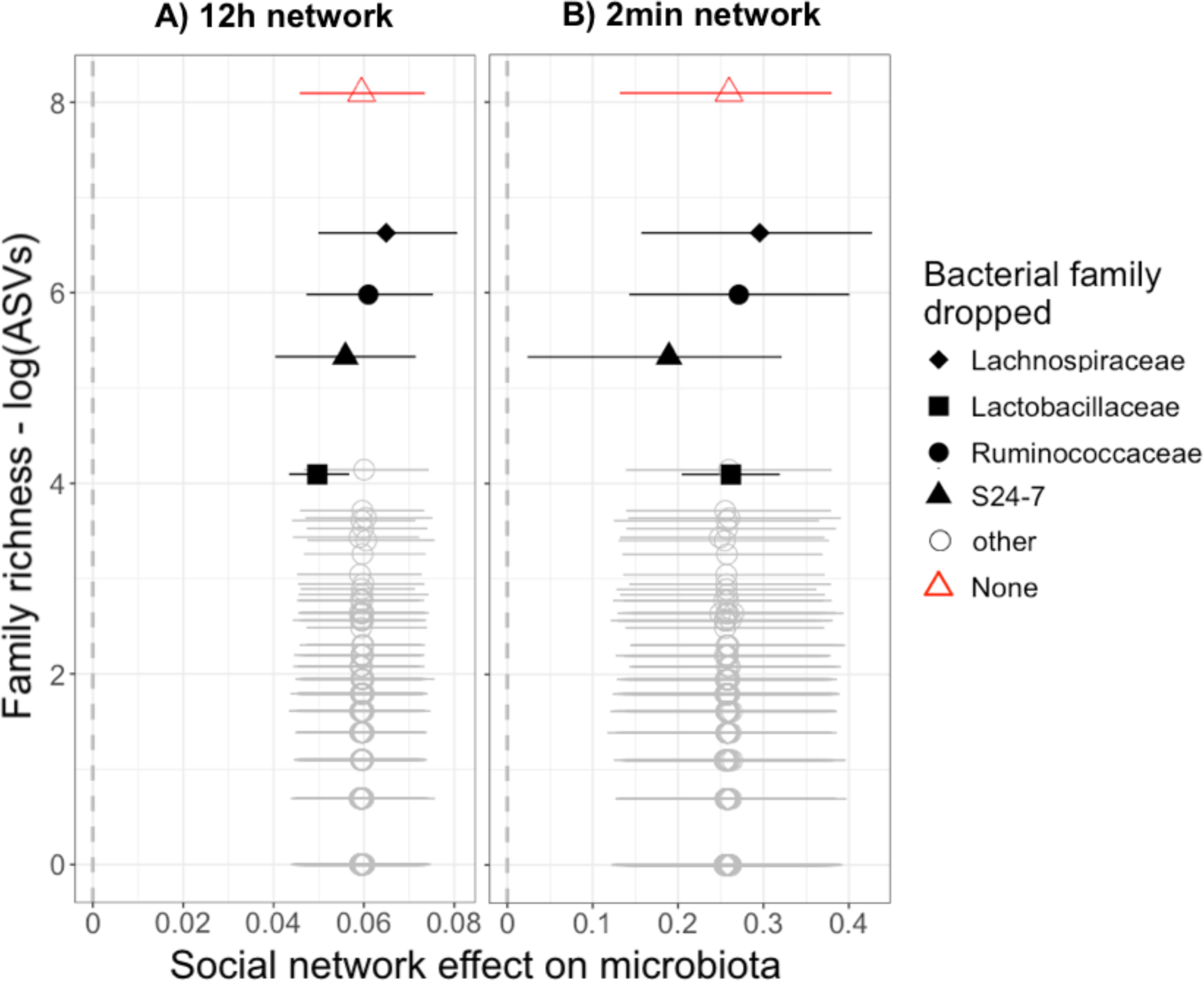
The influence of specific bacterial families on social network effect size. Social network effect sizes (slope of the relationship between social association strength and microbiota similarity, Jaccard Index) and 95% credible intervals are plotted from 146 Bayesian regression (MCMCglmm) models, in which a single bacterial family was excluded from the calculation of microbiota similarity. Effects are plotted against the species richness of each dropped family (logged number of ASVs, y-axis). Results are shown from models using A) the least intimate, 12-hour network and B) the most intimate, 2-minute network.

## Discussion

Recent studies have shown that the social environment can strongly affect gut microbiota composition in group-living species, such as primates living in large groups (26,29) or smaller family units (27,28,30). Here, we provide the first evidence for similar effects in a non-group-living species. The social network of wood mice showed no clear clustering, as those of group-living species do. Yet, the social network strongly predicted similarity among individuals in gut microbiota composition, and this effect was far stronger than effects of spatial or temporal proximity, kinship, and similarity in other host attributes (age, sex). In short, mice who were observed at the same location within the same short timeframe, shared more gut bacterial taxa than mice who were observed together less often. This social effect was sufficiently strong that mice who were observed together even once shared more bacterial taxa than mice who were never observed together.

Social environment effects on the microbiota can result from social partners having more similar environmental exposures, and previous studies have struggled to separate such influences from the effect of social transmission. Here, several findings suggest the social effect we see is likely driven by social transmission, rather than shared exposures. First, we find a strong social network effect even when controlling for host spatial and temporal proximity as well as kinship, reducing the likelihood it is driven by shared traits or exposure to microbes from common environmental sources, such as soil. Second, more intimate definitions of social association (mice co-occurring within a two-minute period, rather than simply during the same night) predicted microbiota similarity more strongly, suggesting close interaction between hosts is important in driving the effect. Finally, the strength of the social network effect varied according to which bacterial families were included in the analysis, in ways that are consistent with a social transmission explanation. When members of the anaerobic, non-spore-forming bacterial family S24-7 (Bacteroidales, Muribaculaceae; 55) were excluded, the social network effect weakened. Conversely, when members of the spore-forming family Lachnospiraceae were excluded (which are able to survive outside the host and have been found in soils; 56,57), the social network effect became slightly stronger. These observations suggest that microbial transmission during close host contact is an important driver of the social effect, allowing hosts to share microbes that cannot persist in the external environment. Previous work in hominids has also shown high host fidelity and even cospeciation with the host among members of the Bacteroidales, while Lachnospiraceae members showed low host fidelity and frequent host switches (58). Taken together, these findings are consistent with the idea that microbes unable to persist outside the host are more reliant on transmission by close contact (e.g. social behaviour or birth), and perhaps in part because of this, they may evolve increased host specificity.

Our findings also indicate that the link between social interactions and the gut microbiota might be more nuanced than previously thought. We found that the strength of social influence on the microbiota varied according to the sex of interacting partners, with social association strength predicting microbiota similarity for male-female and male-male pairs, but not significantly so for female-female pairs. This suggests that behaviours which vary in type, frequency or strength according to the sex of social partners, are involved in gut microbial transmission. In wood mice, home range overlap is much greater among male-female and male-male dyads than among female-female dyads (39,59) and limited data also implies that co-nesting may be more common among male-female pairs than among same-sex pairs (40). Female wood mice are therefore expected to socially interact with one another less often, and female-female links in our social networks may reflect actual social contact to a lesser extent than male-female and male-male links. In line with our findings, recent work found that interactions involving males were more important for the transmission of a herpesvirus pathogen in wood mice (60), potentially suggesting the spread of infectious agents more broadly may be more male-driven in this species. Our findings seem to constitute a mirror image of the common trend in primates, where female-female social bonds are often physically closer than male-male bonds (61), and where social interactions among females have been shown to predict microbiota similarity more strongly than those among males (62,63). In pair-bonding species like humans, the strongest microbiota-homogenizing effects of social interaction may occur in close sexual relationships (37). Interestingly, in wood mice (which do not pair-bond), we find no evidence that male-female associations predict microbiota similarity more strongly than male-male associations. This might be because sexual relationships are not well-captured by our measure of social association, or because other social behaviours prevalent among males are more important in transmission of gut microbes than behaviours specific to mixed-sex pairs.

In addition to social contact homogenising the gut microbiota, we also found that the diversity of an individual’s microbiota is predicted by their position in the social network. Individuals with a central position in the social network, particularly with many contacts or in positions that bridged different parts of the network, carried more bacterial taxa in their gut. Of all network metrics, the strongest predictor of microbiota richness was the number of others an individual was connected to in the network (degree). Similar trends were previously reported in sifakas (28) and chimpanzees (29), and humans self-reporting more social relationships also had greater gut microbial diversity (63). However, effects in the opposite direction have also been found. In barn-swallows, the extent of same-sex social interaction was negatively correlated with microbiota diversity (64) and in red-bellied lemurs, the most sociable individuals had the lowest gut microbiota diversity (27). Perhaps a more careful consideration of social connectedness patterns may help in understanding how sociability might shape microbiota diversity. For example, the sheer amount of social interaction (definition of sociability in 27) might be less important in diversifying the microbiota than the number of transmission sources (definition of sociability in 28). We find that social interactions predict both alpha- and beta-diversity of the gut microbiota – social network position predicted community richness, and social partners had more similar community compositions. Metacommunity theory predicts that connectivity among local communities (hosts) is critical to explaining overall patterns of diversity. On average, dispersal (microbial transmission through host social interaction) is expected to diversify local communities up to a point, by providing novel colonists and rescuing rare species from extinction, but then cease to be enriching as high dispersal begins to homogenize communities and the best competitors at a regional scale come to dominate and exclude others (65). In other words, local diversity is expected to be maximal at intermediate average levels of dispersal (66). If social connectivity is uneven among hosts (as is common in social networks, including ours), a metacommunity could also maintain both diversifying flux and a level of local community uniqueness, that allows competing microbial species to coexist within the metacommunity. In such a network, hosts that interact with many others, especially those likely to harbour distinct microbes, may experience the most diversifying effects of social transmission, compared to those interacting with the same or similar individuals. Consistent with this idea, we found that hosts interacting with others from different parts of the network (with high bridge-type centrality) had more diverse microbiotas, while this was not true for highly connected individuals with more interconnected partners (i.e. with high eigenvector centrality).

Overall, our findings suggest the social environment is an underestimated force shaping the gut microbiota among free-living animals. An important future question then is what role this “social microbiome” (21) plays in host fitness. Besides the pathogenic challenges arising from social contact, which have been acknowledged for some time (67–69) there may also be benefits. Our results suggest social transmission affects microbiota attributes that have potential relevance for host health: microbiota diversity, similarity among interacting individuals, and transmission of anaerobes. While exact relationships between microbiota diversity and beneficial functions remain poorly understood (70), a diverse microbiota might bring benefits in terms of both resisting pathogenic infection (11,71) and increasing metabolic capacity (1,72). Immunological benefits may also result from microbiota similarity among closely interacting individuals. Since symbiotic microbes can be pathogenic in an unaccustomed individual (71,72), sharing a set of familiar microbes with social partners might help maintaining diversity, while minimizing the threat of opportunist pathogens (20,73). Lastly, if anaerobic, non-spore-forming microbes are less likely to be harmful (22) and more likely to be beneficial, social interactions may facilitate the sharing of functionally important, and perhaps more host-specialist symbionts, such as members of the Bacteroidales (58,76). Since such benefits of social behaviour could be present even without any others (e.g. benefits of cooperative behaviour), social transmission of gut microbes could represent an underappreciated force in the early evolution of sociality.

## Acknowledgements

We thank Aurelio Malo for loan of the PIT-tag loggers and advice on field experimental design, Saskia Ricks, Terence Chung, Alice Weightman, Rohan Raval and Joe Williamson for assistance with fieldwork, and Mike Francis for manufacture of loggers and support with their use. We also thank Paul Buerkner for help with brms models, Kevin Foster for helpful comments on the manuscript, and Kirsty Marsh for insightful discussions. This work was funded by a NERC independent Research Fellowship to SCLK (NE/L011867/1), and a Clarendon Scholarship to AR. JAF was supported by a research fellowship from Merton College and BBSRC (BB/S009752/1) and acknowledges funding from NERC (NE/S010335/1). AH acknowledges funding from University of Helsinki for genotyping expenses.

## Contributions

SCLK conceived and designed the study, BA and SCLK carried out fieldwork, collected samples and cleaned field data for analysis, AR conducted microbiome laboratory work, analysed the data and led writing of the manuscript. TT and AH performed mouse genotyping and built the pedigree, JF helped with social network analysis, TC and SCLK provided guidance during analyses, and all authors contributed to and reviewed the manuscript.

## References

1. Turnbaugh PJ, Ley RE, Mahowald MA, Magrini V, Mardis ER, Gordon JI. An obesity-associated gut microbiome with increased capacity for energy harvest. Nature. 2006; 444(7122):1027–1031.

2. Monachese M, Burton JP, Reid G. Bioremediation and Tolerance of Humans to Heavy Metals through Microbial Processes: a Potential Role for Probiotics? Downloaded from. Appl Environ Microbiol. 2012; 78:6397–6404.

3. Chevalier C, Stojanovi O, Colin DJ, Suarez-Zamorano N, Tarallo V, Veyrat-Durebex C et al. Gut Microbiota Orchestrates Energy Homeostasis during Cold. Cell. 2015; 163: 1360–1374.

4. Kamada N, Kim Y-G, Sham HP, Vallance BA, Puente JL, Martens EC, et al. Regulated virulence controls the ability of a pathogen to compete with the gut microbiota. Science. 2012; 336(6086):1325–1329.

5. Zhang N, He Q-S. Commensal Microbiome Promotes Resistance to Local and Systemic Infections. Chin Med J. (Engl) 2015; 128(16):2250–2255.

6. Hooper L V, Littman DR, Macpherson AJ. Interactions Between the Microbiota and the Immune System. Science. 2012; 336(6086):1268–1273.

7. Honda K, Littman DR. The microbiota in adaptive immune homeostasis and disease. Nature. 2016; 535(7610):75–84.

8. Thaiss CA, Zmora N, Levy M, Elinav E. The microbiome and innate immunity. Nature 2016; 535(7610):65–74.

9. Hanski I, von Hertzen L, Fyhrquist N, Koskinen K, Torppa K, Laatikainen T, et al. Environmental biodiversity, human microbiota, and allergy are interrelated. Proc Natl Acad Sci USA. 2012; 109(21):8334–8339.

10. De Luca F, Shoenfeld Y. The microbiome in autoimmune diseases. Clin Exp Immuno.l 2019; 195(1):74–85.

11. Costello EK, Stagaman K, Dethlefsen L, Bohannan BJM, Relman DA. The application of ecological theory toward an understanding of the human microbiome. Science. 2012; 336(6086):1255–1262.

12. Dominguez-Bello MG, Costello EK, Contreras M, Magris M, Hidalgo G, Fierer N, et al. Delivery mode shapes the acquisition and structure of the initial microbiota across multiple body habitats in newborns. Proc Natl Acad Sci USA. 2010; 107(26):11971–11975.

13. Ferretti P, Pasolli E, Tett A, Asnicar F, Gorfer V, Fedi S, et al. Mother-to-Infant Microbial Transmission from Different Body Sites Shapes the Developing Infant Gut Microbiome. Cell Host Microbe. 2018; 24(1):133–145.

14. Lane AA, McGuire MK, McGuire MA, Williams JE, Lackey KA, Hagen EH, et al. Household composition and the infant fecal microbiome: The INSPIRE study. Am J Phys Anthropol. 2019; 169(3):526–539

15. Lehtimäki J, Karkman A, Laatikainen T, Paalanen L, von Hertzen L, Haahtela T, et al. Patterns in the skin microbiota differ in children and teenagers between rural and urban environments. Sci Rep. 2017; 7(1):45651.

16. Tamburini S, Shen N, Wu H, medicine JC-N, 2016 undefined. The microbiome in early life: implications for health outcomes. Nature medicine. 2016; 22(7):713–722.

17. Turnbaugh PJ, Ridaura VK, Faith JJ, Rey FE, Knight R, Gordon JI. The Effect of Diet on the Human Gut Microbiome: A Metagenomic Analysis in Humanized Gnotobiotic Mice. Sci Transl Med. 2009; 1(6):6ra14.

18. David LA, Maurice CF, Carmody RN, Gootenberg DB, Button JE, Wolfe BE, et al. Diet rapidly and reproducibly alters the human gut microbiome. Nature. 2014; 505(7484):559–563.

19. Ottman N, Ruokolainen L, Suomalainen A, Sinkko H, Karisola P, Lehtimäki J, et al. Soil exposure modifies the gut microbiota and supports immune tolerance in a mouse model. J Allergy Clin Immunol. 2019; 143(3):1198–1206.

20. Grieneisen LE, Charpentier MJE, Alberts SC, Blekhman R, Bradburd G, Tung J, et al. Genes, geology and germs: gut microbiota across a primate hybrid zone are explained by site soil properties, not host species. Proc R Soc B Biol Sci. 2019; 286(1901):20190431.

21. Sarkar A, Harty S, Johnson KV-A, Moeller AH, Archie EA, Schell LD, et al. Microbial transmission in animal social networks and the social microbiome. Nat Ecol Evol. 2020; 4(8):1020–1035.

22. Moeller AH, Suzuki TA, Phifer-Rixey M, Nachman MW. Transmission modes of the mammalian gut microbiota. Science. 2018; 362(6413):453–457.

23. Hufeldt MR, Nielsen DS, Vogensen FK, Midtvedt T, Hansen AK. Variation in the gut microbiota of laboratory mice is related to both genetic and environmental factors. Comparative medicine. 2010; 60(5):336–347.

24. Hildebrand F, Nguyen TLA, Brinkman B, Yunta R, Cauwe B, Vandenabeele P, et al. Inflammation-associated enterotypes, host genotype, cage and inter-individual effects drive gut microbiota variation in common laboratory mice. Genome Biol. 2013; 14(1):R4.

25. Lees H, Swann J, Poucher SM, Nicholson JK, Holmes E, Wilson ID, Marchesi JR. Age and microenvironment outweigh genetic influence on the Zucker rat microbiome. PloS one. 2014; 9(9):e100916.

26. Tung J, Barreiro LB, Burns MB, Grenier JC, Lynch J, Grieneisen LE, et al. Social networks predict gut microbiome composition in wild baboons. Elife. 2015; 4:e05224.

27. Raulo A, Ruokolainen L, Lane A, Amato K, Knight R, Leigh S, et al. Social behaviour and gut microbiota in red-bellied lemurs (Eulemur rubriventer): In search of the role of immunity in the evolution of sociality. J Anim Ecol. 2018; 87(2):388–399.

28. Perofsky AC, Lewis RJ, Abondano LA, Di Fiore A, Meyers LA. Hierarchical social networks shape gut microbial composition in wild Verreaux’s sifaka. Proceedings Biol Sci. 2017; 284(1868):20172274

29. Moeller AH, Foerster S, Wilson ML, Pusey AE, Hahn BH, Ochman H. Social behavior shapes the chimpanzee pan-microbiome. Sci Adv. 2016; 2(1):e1500997.

30. Wikberg EC, Christie D, Sicotte P, Ting N. Interactions between social groups of colobus monkeys (Colobus vellerosus) explain similarities in their gut microbiomes. Anim Behav. 2020; 163:17–31.

31. Bennett G, Malone M, Sauther ML, Cuozzo FP, White B, Nelson KE, et al. Host age, social group, and habitat type influence the gut microbiota of wild ring-tailed lemurs (Lemur catta). Am J Primatol. 2016; 78(8):883–892.

32. Theis KR, Schmidt TM, Holekamp KE. Evidence for a bacterial mechanism for group-specific social odors among hyenas. Sci Rep. 2012; 2:615 .

33. Leclaire S, Nielsen JF, Drea CM. Bacterial communities in meerkat anal scent secretions vary with host sex, age, and group membership. Behav Ecol. 2014;25(4):996–1004.

34. Antwis RE, Lea JMD, Unwin B, Shultz S. Gut microbiome composition is associated with spatial structuring and social interactions in semi-feral Welsh Mountain ponies. Microbiome. 2018; 6(1):207.

35. Song SJ, Lauber C, Costello EK, Lozupone CA, Humphrey G, Berg-Lyons D, et al. Cohabiting family members share microbiota with one another and with their dogs. Elife. 2013;2:e00458

36. Grieneisen LE, Livermore J, Alberts S, Tung J, Archie EA. Group Living and Male Dispersal Predict the Core Gut Microbiome in Wild Baboons. Integr Comp Biol. 2017; 57(4):770–785.

37. Dill-McFarland KA, Tang Z-Z, Kemis JH, Kerby RL, Chen G, Palloni A, et al. Close social relationships correlate with human gut microbiota composition. Sci Rep. 2019; 9(1):703.

38. Wilson EO. Elementary concepts in sociobiology. In: Wilson EO. Sociobiology: The New Synthesis. 25^th^ edn. (Harvard University Press, Cambridge, Massachutes, USA, 2000) p. 8.

39. Godsall B. Mechanisms of space use in the wood mouse, Apodemus sylvaticus. Doctoral Thesis, Imperial College, London 2015.

40. Wolton RJ. The ranging and nesting behaviour of Wood mice, Apodemus sylvaticus (Rodentia: Muridae), as revealed by radio-tracking. J Zool. 2009; 206(2):203–222.

41. Godsall B, Coulson T, Malo AF. From physiology to space use: energy reserves and androgenization explain home-range size variation in a woodland rodent. J Anim Ecol. 2014; 83(1):126–135.

42. Wang J, Santure AW. Parentage and sibship inference from multilocus genotype data under polygamy. Genetics. 2009; 181(4):1579–1594.

43. Farine DR. Animal social network inference and permutations for ecologists in R using asnipe. Methods Ecol Evol. 2013; 4(12):1187–1194.

44. Csardi G, Nepusz T. The igraph software package for complex network research. InterJournal, complex systems. 2006; 1695(5):1–9.

45. Firth JA, Sheldon BC. Social carry-over effects underpin trans-seasonally linked structure in a wild bird population. Ecol lett. 2016; 19(11):1324–32.

46. Callahan BJ, McMurdie PJ, Rosen MJ, Han AW, Johnson AJA, Holmes SP. DADA2: High-resolution sample inference from Illumina amplicon data. Nat Methods. 2016; 13(7):581–583.

47. McMurdie PJ, Holmes S. phyloseq: An R Package for Reproducible Interactive Analysis and Graphics of Microbiome Census Data. PLoS One. 2013; 8(4):e61217.

48. McKnight DT, Huerlimann R, Bower DS, Schwarzkopf L, Alford RA, Zenger KR. Methods for normalizing microbiome data: An ecological perspective. Methods Ecol Evol. 2019; 10(3):389–400.

49. Hsieh TC, Ma KH, Chao A. iNEXT: an R package for rarefaction and extrapolation of species diversity (Hill numbers). Methods Ecol Evol. 2016; 7(12):1451–1456.

50. Oksanen J, Kindt R, Legendre P, O’Hara B, Stevens MH, Oksanen MJ, Suggests MA. The vegan package. Community ecology package. 2007; 10(631-637):719.

51. Bürkner PC. brms: An R Package for Bayesian Multilevel Models Using Stan. J Stat Softw. 2017; 80(1):1–28.

52. Bürkner PC. Advanced Bayesian Multilevel Modeling with the R Package brms. The R Journal. 2018; 10(1): 395–411.

53. Hadfield JD. MCMC methods for multi-response generalized linear mixed models: the MCMCglmm R package. J Stat Softw. 2010; 33(2):1–22.

54. Dekker D, Krackhardt D, Snijders TA. Sensitivity of MRQAP tests to collinearity and autocorrelation conditions. Psychometrika. 2007; 72(4):563–581.

55. Ormerod KL, Wood DLA, Lachner N, Gellatly SL, Daly JN, Parsons JD, et al. Genomic characterization of the uncultured Bacteroidales family S24-7 inhabiting the guts of homeothermic animals. Microbiome. 2016; 4(1):36.

56. Wieczorek AS, Schmidt O, Chatzinotas A, von Bergen M, Gorissen A, Kolb S. Ecological Functions of Agricultural Soil Bacteria and Microeukaryotes in Chitin Degradation: A Case Study. Front Microbiol. 2019; 10:1293.

57. Huang X, Liu L, Zhao J, Zhang J, Cai Z. The families Ruminococcaceae, Lachnospiraceae, and Clostridiaceae are the dominant bacterial groups during reductive soil disinfestation with incorporated plant residues. Applied Soil Ecology. 201; 135:65–72.

58. Moeller AH, Caro-Quintero A, Mjungu D, Georgiev A V, Lonsdorf E V, Muller MN, et al. Cospeciation of gut microbiota with hominids. Science. 2016; 353(6297):380–382.

59. Tew TE, Macdonald DW. Dynamics of space use and male vigour amongst wood mice, Apodemus sylvaticus, in the cereal ecosystem. Behav Ecol Sociobiol 1994 May; 34(5):337–45.

60. Erazo D, Pedersen AB, Gallagher K, Fenton A. Who acquires infection from whom? Estimating herpesvirus transmission rates between wild rodent host groups. bioRxiv 2020; 2020.09.18.302489.

61. Taylor SE, Klein LC, Lewis BP, Gruenewald TL, Gurung RAR, Updegraff JA. Biobehavioral responses to stress in females: Tend-and-befriend, not fight-or-flight. Psychol Rev. 2000; 107(3):411–29.

62. Amato KR, Van Belle S, Di Fiore A, Estrada A, Stumpf R, White B, et al. Patterns in Gut Microbiota Similarity Associated with Degree of Sociality among Sex Classes of a Neotropical Primate. Microb Ecol. 2017; 74(1):250–258.

63. Brito IL, Gurry T, Zhao S, Huang K, Young SK, Shea TP, et al. Transmission of human-associated microbiota along family and social networks. Nat Microbiol. 2019; 4(6):964–971.

64. Johnson KV-A. Gut microbiome composition and diversity are related to human personality traits. Hum Microbiome J. 2020; 15:100069.

65. Levin II, Zonana DM, Fosdick BK, Song SJ, Knight R, Safran RJ. Stress response, gut microbial diversity and sexual signals correlate with social interactions. Biol Lett. 2016; 12(6):20160352.

66. Mouquet N, Loreau M. Community Patterns in Source-Sink Metacommunities. Am Nat. 2003; 162(5):544–557.

67. Altizer S, Nunn CL, Thrall PH, Gittleman JL, Antonovics J, Cunningham AA, et al. Social organization and parasite risk in mammals: Integrating theory and empirical studies. Annu Rev Ecol Evol Syst. 2003; 34:517–547.

68. Loehle C. Social Barriers to Pathogen Transmission in Wild Animal Populations. Ecology. 1995; 76(2):326–335.

69. Moeller A, Dufva R, Allander K. Parasites and the Evolution of Host Social Behavior. Adv Study Behav. 1993; 22:65–102.

70. Reese AT, Dunn RR. Drivers of microbiome biodiversity: A review of general rules, feces, and ignorance. MBio. 2018; 9(4).

71. Pallen MJ. The human microbiome and host-pathogen interactions. In: Metagenomics of the Human Body. Springer New York; 2011; p. 43–61.

72. Amato KR, Leigh SR, Kent A, Mackie RI, Yeoman CJ, Stumpf RM, et al. The Gut Microbiota Appears to Compensate for Seasonal Diet Variation in the Wild Black Howler Monkey (Alouatta pigra). Microb Ecol. 2014; 69(2):434–443.

73. Barribeau SM, Villinger J, Waldman B. Ecological immunogenetics of life-history traits in a model amphibian. Biol Lett. 2012; 8(3):405–407.

74. Feng T, Elson CO. Adaptive immunity in the host-microbiota dialog. Mucosal Immunol. 2011; 4(1):15–21.

75. Amato KR. Incorporating the Gut Microbiota Into Models of Human and Non-Human Primate Ecology and Evolution. Am J Phys Anthropol. 2016; 159:196–215 .

76. Knowles SCL, Eccles RM, Baltrūnaitė L. Species identity dominates over environment in shaping the microbiota of small mammals. Ecol Lett. 2019; 22(5):826–837.

