## Supplementary Material for "Social networks strongly predict the gut microbiota of wild mice"

**Index**

**Supplementary Materials and Methods**

1. Additional trapping and sampling methods
2. Behavioural data collection using PIT-tag loggers
3. Mouse genotyping and pedigree reconstruction
4. Gut microbiota analysis using 16S rRNA profiling
5. Alternative statistical frameworks for predicting microbiota composition with social association strength
6. Null models describing the effect of reduced network density on estimates of models predicting microbiota with social association

**Supplementary Figures**

**Figure S1****:** Logger study design and logger effort across the grid

**Figure S2****:** Custom PIT-tag logger

**Figure S3****:** Logger data filtering

**Figure S4****:** Microbial standard mock community profile

**Figure S5****:** iNEXT analysis results

**Figure S6****:** Distribution of bacterial taxon (ASV) relative abundances across all samples

**Figure S7****:** Distribution of microbiota similarity metrics

**Figure S8****:** Temporal dynamics of the wood mouse microbiota

**Figure S9**: MRQAP model estimates for the effect of spatial distance on social association strength across networks

**Figure S10**: No single dyad drives the effect of social network on microbiota similarity

**Supplementary Tables**

**Table S1**: PCR protocols and multiplexing for 12 microsatellite loci used to genotype wild-caught wood mice

**Table S2**: Summary data for the final 11 microsatellite loci used in pedigree reconstruction.

**Table S3**: The relative effects of individual identity, temporal change and methodological factors on gut microbiota compositional variation

**Table S4**: The extent of individual-level variation (repeatability) in gut microbiota after controlling for known individual-level covariates

**Table S5****:** The effect of demographic individual-level variables on gut microbiota composition.

**Table S6****:** Correlations among social networks with varying edge definitions

**Table S7**: Variables shaping social association strength (Adjusted SRI) across networks

**Table S8****:** Results of *brms* models testing the effect of social association strength and covariates on microbiota similarity (Jaccard Index)

**Table S9****:** Results of MCMCglmm models testing the effect of social association strength and covariates on microbiota similarity

**Table S10****:** Results of MRQAP models testing the effect of social association strength and covariates on microbiota similarity (Jaccard Index)

**Table S11**: Results of *brms* models testing the effect of binary social association (BI) and covariates on microbiota similarity (Jaccard Index)

**Table S12****:** Results of *brms* models testing whether the effect of social association strength on microbiota similarity (Jaccard Index) varies according to the sex of interacting individuals.

**Table S13****:** Results of MCMCglmm models predicting microbiota diversity with covariates

**Table S14****:** Effects of social centrality metrics and non-social factors on gut microbiota Shannon diversity

**Supplementary References**

**Supplementary Materials and Methods**

**Additional trapping and sampling methods**

Data for this study was collected as part of a longer-term rodent capture-mark-recapture study. While several rodent species were caught (*Apodemus sylvaticus*, *Apodemus flavicollis* and *Myodes glareolus*), we focus on the most abundant species, wood mice (*Apodemus sylvaticus*). Trapping was performed every 2-4 weeks, using 122 small folding Sherman traps (5.1 x 6.4 x 16.5cm, H. B Sherman). Traps baited with eight peanuts, a slice of apple and sterile cotton wool for bedding were set at dusk and collected at dawn, with all animals processed, sampled and then released inside the 100m^2^ grid cell they were captured in. As part of processing, captured individuals were identified to species, sexed, weighed, and aged (juvenile or adult) based on size and pelage characteristics. Ear punches were collected from all mice and stored in ethanol at -20°C to provide genetic material for host genotyping. Faecal samples for gut microbiota analysis were collected from the trap and frozen at -80^o^C within 8 hours of collection. Utensils used for faecal pellet collection (e.g. tweezers) were sterilized using 70% ethanol between each sample processed, and all traps showing evidence of rodent contact were washed in bleach solution and autoclaved between trapping sessions.

**Behavioural data collection using PIT-tag loggers**

PIT-tag loggers took the form of a large plastic box with two entrance tubes (Figure S2A) leading to a central wooden box containing sawdust and a single peanut (Figure S2B). The peanut was used to give a sense of the frequency with which untagged animals visited the loggers. An RFID coil surrounding the entrance to the central box recorded the PIT-tag ID of any tagged rodent present under it every 0.3 seconds. Peanut oil around the tube entrances, renewed at each logger rotation, was used as a minor lure. Loggers were powered using 12V lead acid rechargeable batteries. Further details about the loggers can be found in (1).

Initially 9 loggers were distributed across the plot, though for a period of 6 weeks (April-May 2015) one logger was broken and only 8 were used during this time. We designed a logger rotation plan such that each logger had an assigned “territory” (plot region) including a constant number of contiguous 100m^2^ grid cells, and was moved between these in order to achieve even spatial coverage of the plot throughout the study period. With 9 loggers (as used for the vast majority of the study), each logger had a territory of 27 contiguous 100m^2^ grid cells (Figure S1A). At each rotation, an R script was used to select the new locations to which loggers would move: each logger was moved to a new grid cell selected randomly without replacement from those in its territory, until all grid cells had been monitored, before the selection process was repeated. The precise (1m^2^) position of each logger within a grid cell was also randomly selected at each rotation. Loggers were always rotated between 10am and 2pm when wood mice (which are strongly nocturnal) are least active. Rotations were performed on average 3.54 times per week throughout the year (range 2-7 times per week). When loggers were left for several consecutive nights in a location for logistical reasons (e.g. over a weekend), only data from the first of these nights was used in subsequent analyses, to maintain even spatial coverage in the final dataset (Figure S1B). Loggers were deliberately not moved to a new location just after a trapping night, as logger data from trapping nights (when many mice may be in traps and therefore not recorded) was also intentionally filtered out of the data before analysis. Logger data rows containing corrupt tag reads was also removed during data cleaning, prior to analysis (See Figure S3 for details on how much data was lost in each step).

**Mouse genotyping and pedigree reconstruction**

*Genotyping*

Genomic DNA was extracted from tissue samples (ear clips) using QIAamp DNA Micro Kits (Qiagen). Twelve microsatellite loci were selected for *Apodemus sylvaticus* from the literature (2,3). Target regions were amplified in four multiplex PCRs designed according to target region size range and annealing temperature (Table S1). PCR conditions were as follows: initial denaturation at 95 °C for 15min, 30 cycles involving 95 °C for 30s, the annealing temperature (Table S1) for 90s, and extension at 72 °C for 1min, followed by a final extension at 60 °C for 10min. PCR reaction volume was 10μl containing 1μl template DNA, 0.5μl each primer (at 10 μmol concentration), 5μl Qiagen Multiplex PCR Master Mix , 1μl Q-solution and 2μl PCR-grade water. Each PCR plate included one negative (H_2_0) control which was run on a 2% agarose gel to verify lack of contamination. PCR products were prepared for sequencing by diluting to equal concentration (measured using a Qubit fluorometer) and mixing with a solution of deionized 95-100% formamide and size standard (Genescan 500 ROX) before sequencing on a 3730 DNA Analyzer (Applied Biosystems) and scoring in GeneMapper Software (version 5). Genotyping results were evaluated with Cervus 3.0.7 (4) and Micro-checker 2.2 (5), after which one microsatellite marker (GACAB3A) was discarded as it showed significant deviation from Hardy-Weinberg Equilibrium. Summary characteristics for the remaining 11 loci used for pedigree reconstruction are shown in Table S2.

*Pedigree reconstruction*

A pedigree was constructed using COLONY 2.0.6.5 (6) a program for parental and sibship inference from genotype data. This software was chosen because it can perform analyses for polygamous species and accounts for genotyping error. It divides samples into family clusters, in which individuals are related either via sibship or shared parentage. The likelihood of a cluster is calculated based on Mendelian inheritance rules (7). COLONY was run multiple times to get the most accurate estimates for sibship/parentage, adjusting parameter expectations to take into account that wood mice are polygamous (half-sibs are common) and may inbreed (genotype frequencies may be biased towards homozygosity). We compared kinship results from COLONY with trapping data, aided with visualization with Pedigree Viewer to identify impossible relationships (based on age and time trapped). We found 11 impossible mother-pup and 4 impossible father-pup pairs, which were excluded after which the model was re-run, this time estimating kinship without any conflicts with the trapping data. The resulting pedigree contained 17 mother-pup pairs, 14 father-pup pairs, 13 full-sibling-pairs and 26 half-sibling-pairs. This pedigree was used to create dyadic numeric kinship matrices, where kinship was transformed into a numeric distance variable, with for example parent-offspring pairs and full siblings both assigned a kinship value of 0.5 and half-siblings/cousins 0.125.

**Gut microbiota analysis using 16S rRNA profiling**

*Library preparation and sequencing*

DNA was extracted from samples using Zymo Quick-DNA™ Fecal/Soil Microbe 96 kits (Zymo) according to manufacturer's instructions. For extractions as well as subsequent PCRs, samples were randomized across batches/plates to avoid confounding biological and technical effects on downstream microbial data. An approximately 240bp V4 region of the 16S rRNA gene was amplified using primers N515F and N806R (8) in two-step (tailed-tag) approach with dual-indexing (9). In addition to samples, we included one extraction control (PCR-grade water subjected to the DNA extraction method) and one PCR control (PCR-grade water) per 96-well plate as well as 2 extractions of a mock community (ZYMOBiomics Microbial Community Standard cat. No. D6300) to evaluate extraction, PCR and sequencing accuracy.

For all samples a test first round PCR was performed, to check sample amplification and decide on an appropriate number of PCR cycles for the final library preparation. For this test PCR, the following were combined in a 20µl reaction volume: 10µl KAPA 2x Mastermix (KAPA Biosystems), 0.25µl each primer at 10µM, 4.5µl PCR-grade water and 5µl undiluted DNA extraction. Cycling conditions were as follows: denaturation at 98°C for 2min, 35 cycles of 95°C for 20s, 65°C for 15s, 70°C for 30s, followed by a final extension at 70°C for 5min and a 4°C hold. Products were visualized on 2% agarose gels, and no amplification was observed in extraction or PCR controls. Subsequently the first round PCR for use in subsequent sequencing was performed, using exactly the same conditions as described above but with fewer cycles (20 instead of 35).

Unpurified first round PCR products were sent to the Centre for Genomic Research (Liverpool, UK) for indexing PCRs, purification, pooling, amplicon size selection and sequencing. First round PCR products were purified using Agencourt Ampure XP beads (Beckman Coulter, Brea, CA, USA), by adding 20µl beads at room temperature to samples, before washing twice with 200µl 80% ethanol and eluting in 10µl H_2_0. For the second round PCR (addition of indices), the following were combined in a 20µl reaction volume: 10µl KAPA 2x Mastermix (KAPA Biosystems), 0.5µl of each primer at 10µM and 9µl of clean PCR product. For indexing, eight forward primers (i5) and twelve reverse (i7) primers were used, each containing a unique barcode, creating 96 unique combinations (barcode sequences are reported in the Illumina Nextera Protocol (10). Second round PCR products were purified using Agencourt Ampure XP beads as described above, but eluted in 20µl H_2_0. Each second round PCR product (library) was analyzed using a 2100 Bioanalyzer (Agilent Technologies, Santa Clara, CA, USA) and up to 96 libraries pooled per sequencing run, at equimolar concentration of fragments in the expected size range. Each pool was size-selected using a Pippin Prep, by excluding fragments outside the expected range. Libraries were sequenced using 2x250bp paired-end sequencing on an Illumina MiSeq, with samples from different extraction batches randomly allocated to four different runs. Visualization of the composition of the mock samples revealed that our laboratory- and bioinformatics pipeline successfully captured the microbial diversity present in mock samples (Figure S4).

*Bioinformatics*

Sequence data were processed through the DADA2 pipeline (version 1.6.0) (11). *cutadapt* (12) was used to determine optimal trimming parameters for removal of leading primer and adapter sequences (27 base pairs from the beginning of each read), and trimming was performed using the trimLeft argument in DADA2. Following visual inspection of sequence quality, low-quality tails were also trimmed (leaving 230 bp for forward and 180 bp for reverse reads). Sequences were dereplicated, and amplicon sequence variants (ASVs) inferred using the DADA2 algorithm. Forward and reverse paired-end reads were merged after which reads that could not be merged, were putative chimeric sequences or were of abnormal length (<238 bp or > 245 bp) were removed from the dataset.

Within the R package *phyloseq* (13) taxonomy was assigned to ASVs using the GreenGenes Database (GreenGenes Database Consortium 13.8). One sample with exceptionally low read count was removed from the dataset (n=300 reads, while read count >10 000 for all other samples). We then used package iNEXT (14) to confirm that sample completeness estimates plateaued and diversity estimates stabilized at read counts around 2500-4000 reads, well below the read counts for all samples in the remaining dataset (Figure S5). We also used iNEXT to derive sample-level asymptotic estimates of microbial richness and Shannon diversity, which are corrected for the modelled effect of read count. After asymptotic correction, Shannon diversity estimates were not significantly predicted by read count but richness estimates were (linear regressions, p<0.01), and thus effect of read count was controlled for in all models predicting richness. Singleton reads were filtered out of the dataset, to avoid residual bias from possible contaminants or sequencing errors, as well as ASVs assigned to taxa known to not be gut bacteria (Cyanobacteria, Xanthomonadales, Mitochondria). Since the distribution of abundances for remaining taxa showed no obvious tail of rare taxa (Figure S6) further abundance filtering was not performed. Finally, ASV counts were normalized by calculating proportional abundance within each sample (15).

**Alternative statistical frameworks for predicting microbiota composition with social association strength**

To check our results were robust to the statistical approach used, we used two alternative statistical methods to verify key results from the model predicting microbiota composition with the 12h social network. First, another Bayesian regression model from the R package MCMCglmm (16) was used with the same data. Model structure was identical to that used in *brms* models, except that, since this package does not allow beta-regression and since the distribution of response variable values was not skewed, the model used a Gaussian link function. As with *brms* models, the MCMCglmm model was run first with Jaccard Index as the response, and then with Bray-Curtis dissimilarity as the response, to assess whether results held using an abundance-weighted community similarity metric. Second, an alternative matrix permutation-based modelling framework was used, to further verify results with a contrasting analytical method. Here we used multiple regression quadratic assignment procedure (MRQAP, 17), to test the relationship between social proximity and gut microbiota similarity while controlling for the same set of covariates as above. MRQAP is a null model-based matrix permutation test commonly applied in social network analyses (e.g. 18,19). MRQAP was run on a dataset including only one microbiota sample per individual, selected at random in each of 100 different iterations. Finally, to confirm no single dyadic interaction disproportionately influenced results, each dyad was dropped one at a time from the MCMCglmm model, and the effect on coefficients and statistical significance examined. Results from these analyses showing a consistent effect of the social network on microbiota composition are presented in Table S9, Table S10 and Figure S10 respectively.

**Null models describing the effect of reduced network density on estimates of models predicting microbiota with social association**

In order to accurately compare social association-microbiota effect sizes across networks, we had to account for decreasing density in networks with more intimate edge definitions (Figure 2), since it is possible that differences in network density alone could influence effect size differences. We therefore compared observed social association effect size estimates with those derived from artificially thinned versions of the least intimate (12h) network. Six artificial versions of the 12h social network were made by stepwise thresholding (converting the weakest edges to zero), to achieve the same percentage of non-zero-links (=density) as each of our real social networks. In this way, we created a set of artificial networks that had the same density as each of our real networks made using edge definitions of varying intimacy, but that were in fact subsets of stronger links from the 12h network.

**Supplementary Figures**

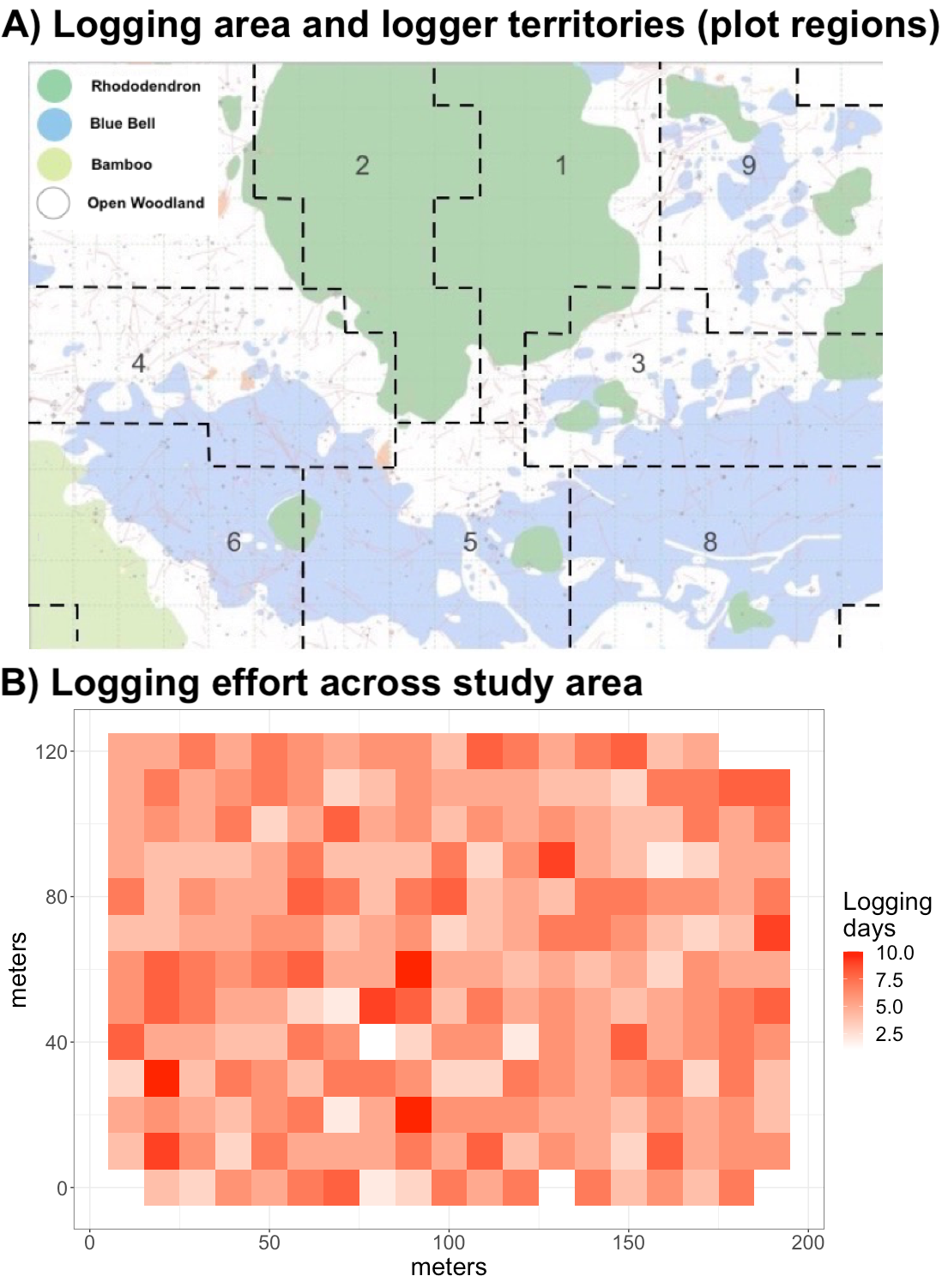

**Figure S1**: **Logger study design and logger effort across the grid**. Map of the 2.43 ha study plot (Nash’s Copse) showing (A) Nine numbered logger territories (plot regions) showing habitat type (rhododendron=dark green, bamboo=light green, open woodland=white or blue) (B) logging effort (total number of logging days performed for each 100m^2^ grid cell).

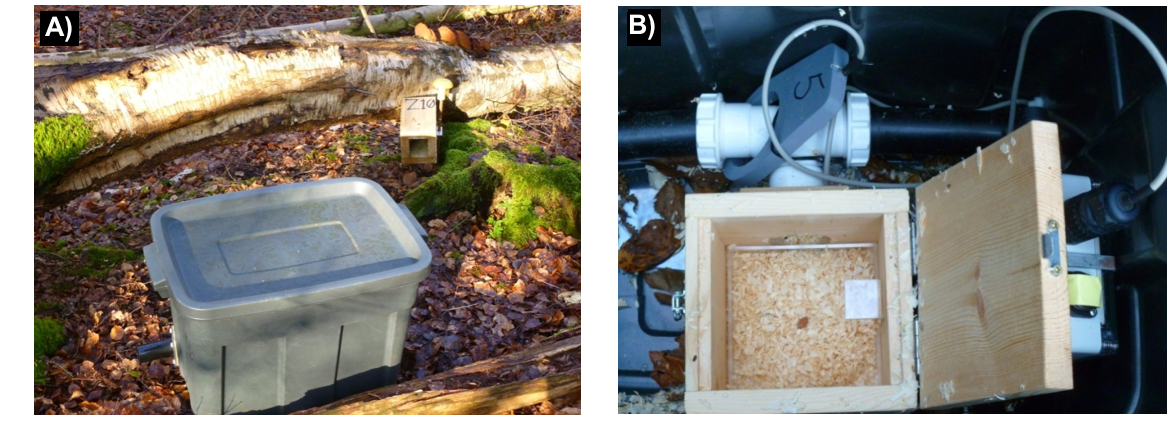

**Figure S2: Custom PIT-tag logger** (A) External view showing two entrance tubes on opposite sides (B) Internal view showing RFID reader (labelled ‘5’) around the entry tubes, and central box containing sawdust and a provisioned with a peanut.

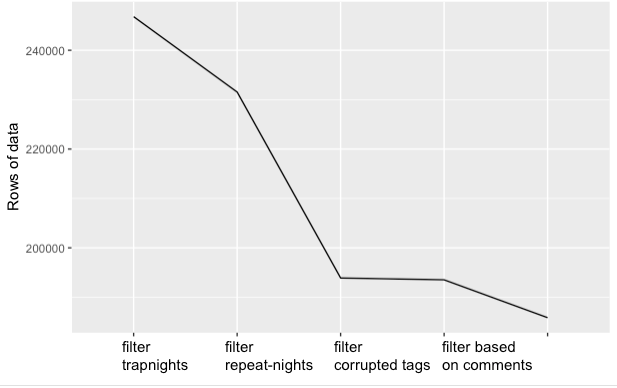

**Figure S3: Logger data filtering.** The effect of each filtering step (x-axis) on the number of logger data rows (y-axis)

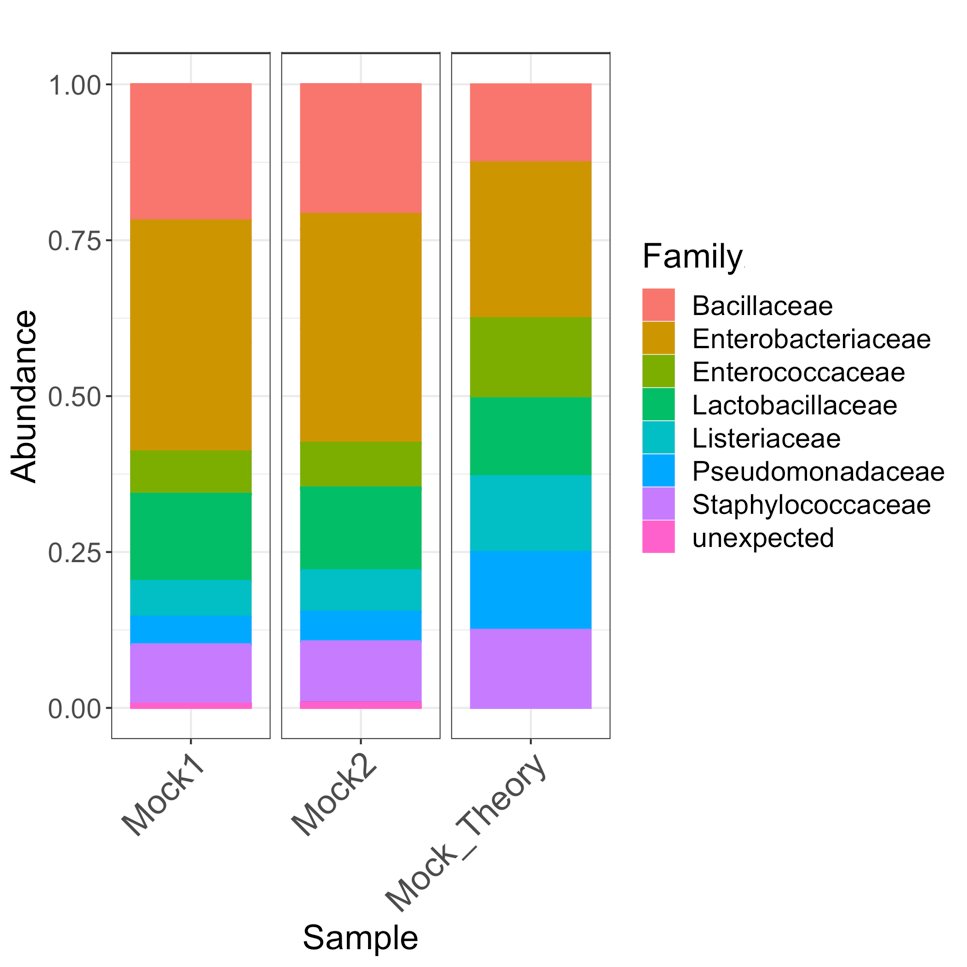

**Figure S4: Microbial standard (mock) community profiles.** Community composition in sequenced mock community aliquots (Mock1, Mock2) compared to the expected composition (Mock_Theory).

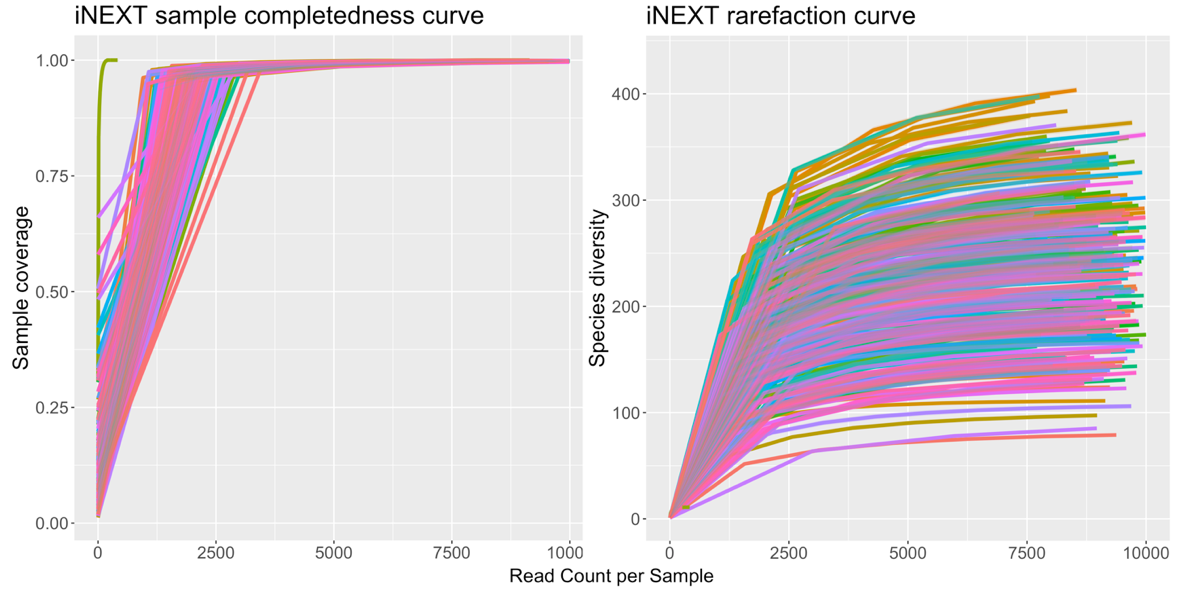

B

A

**Figure S5: iNEXT analysis results** A) Sample completeness curve, showing completeness plateaus above read counts of approximately 4000. B) Rarefaction curve, showing diversity estimates stabilize at read counts above approximately 2500.

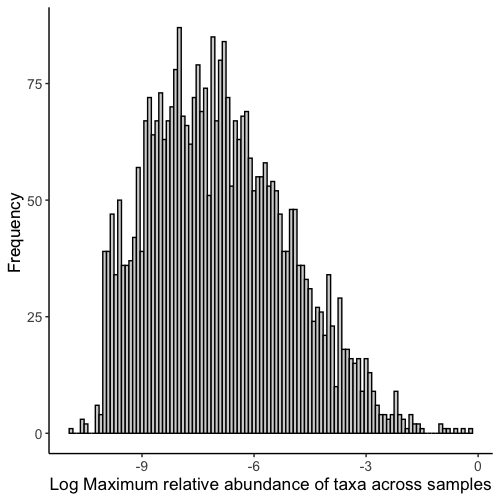

**Figure S6: Distribution of bacterial taxon (ASV) relative abundances across all samples.**

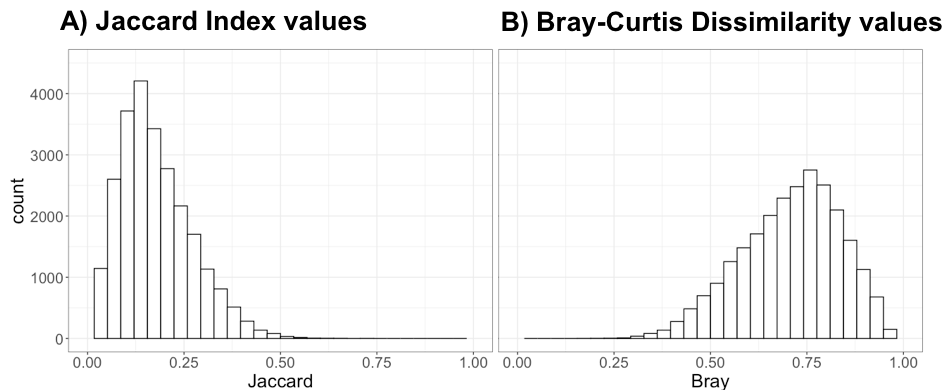

**Figure S7: Distribution of microbiota similarity metrics** A) Jaccard Index of similarity B) Bray-Curtis index of dissimilarity.

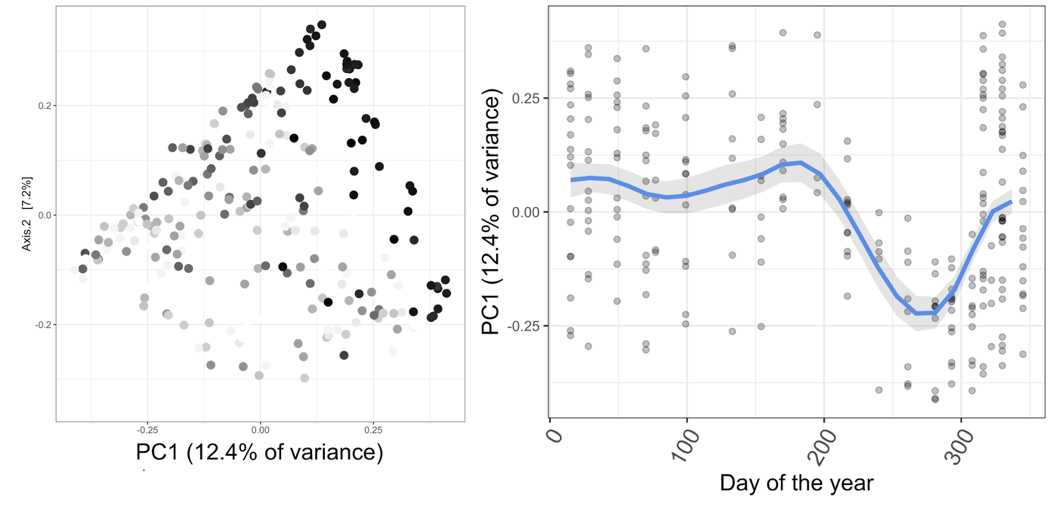

**Figure S8: Temporal dynamics of the wood mouse gut microbiota** A) PCoA on Jaccard distance shows how temporal (seasonal) variation in microbiota composition (darker colour = later in the year) is largely captured by the first axis of variation (PC1). B) PC1 shows marked variation during the year, with a major fluctuation between July and October. A model fit from a generalised additive mixed model (GAMM) is plotted, with the blue line indicating fitted values and shaded grey areas the approximate 95% confidence intervals. The GAMM fits a flexible smoothed function for microbiota variability (PC1 of Jaccard Index) across days of the year.

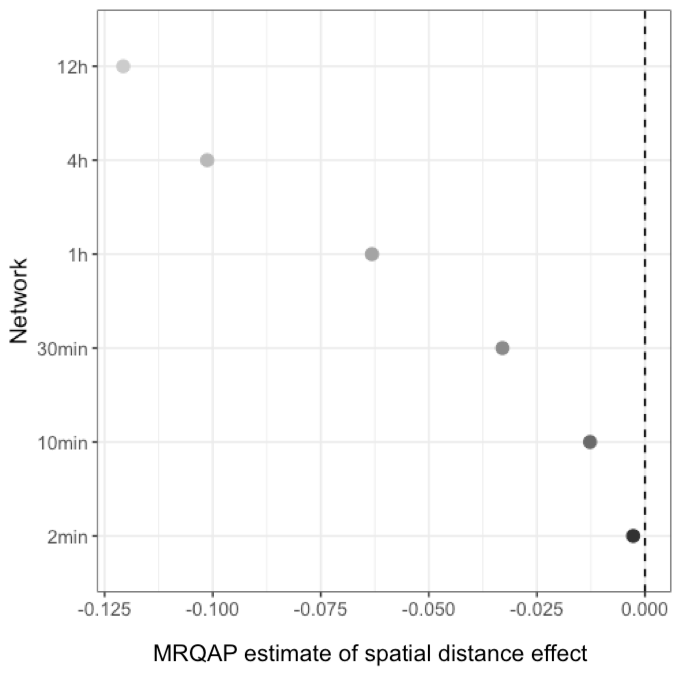

**Figure S9: MRQAP model estimates for the effect of spatial distance on social association strength across networks.** Networks (y-axis, point colour) with increasingly intimate edge definition have lower model estimate predicted by spatial distance. In other words, more intimate social associations are less strongly predicted by spatial proximity between individual mice. Model covariates and results in Table S7.

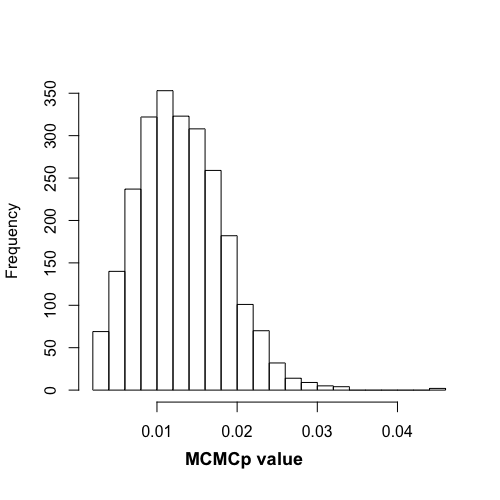

**Figure S10: No single dyad drives the effect of social network on microbiota similarity.** Distribution of p-values for the social network effect from a series of MCMCglmm models that excluded a single dyad from the model. Models used the 12h network to predict microbiota similarity (Jaccard Index). Since p-values remain <0.05 in all models, no single dyad drives the significance of the social network effect.

**Supplementary Tables**

**Table S1**: **PCR protocols and multiplexing for 12 microsatellite loci used to genotype wild-caught wood mice**

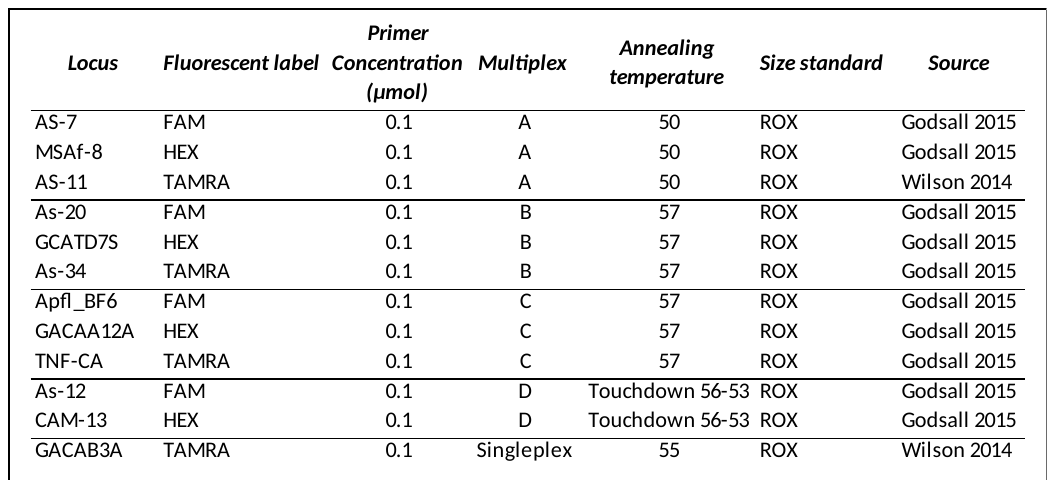

**Table S2: Summary data for the final 11 microsatellite loci used in pedigree reconstruction.** Observed (Hobs) and expected (Hexp) heterozygosity values, deviation from Hardy-Weinberg Equilibrium (HWE, NS = non-significant), null allele and scoring error rates are shown. Null alleles, estimated using COLONY, use the expected distribution of homozygosity under HWE to describe the expected proportion of alleles missed by technical issues, from all alleles in a locus. Scoring error is a measure of total error, due human and technical variation, in repeatability when genotyping the same individual.

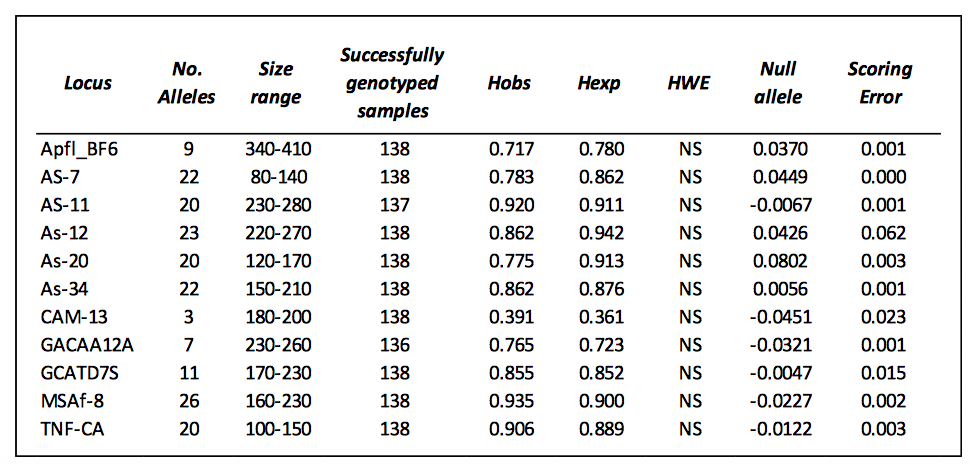

**Table S3: The relative effects of individual identity, temporal change and methodological factors on gut microbiota compositional variation.** Results are from a marginal PERMANOVA on data from mice sampled more than once, with either Jaccard distance or Bray-Curtis dissimilarity as the response. A marginal PERMANOVA was used such that the variance explained by each term does not depend on the order of explanatory variables. Significant terms (p<0.05) are shown in bold.

| **JACCARD** | | | | |
| --- | --- | --- | --- | --- |
|  | **Df** | **F** | **p** | **R^2^** |
| Read count | 1 | 0.956 | 0.526 | 0.003 |
| **PCR plate** | **3** | **1.285** | **0.032** | **0.013** |
| ***Individual ID*** | ***74*** | ***1.321*** | ***0.001*** | ***0.334*** |
| **Month** | **12** | **1.471** | **0.001** | **0.060** |
| Residual | 150 |  |  | 0.512 |
| Total | 240 |  |  | 1.000 |
| **BRAY-CURTIS** | | | | |
|  | **Df** | **F** | **p** | **R^2^** |
| Read count | 1 | 0.993 | 0.437 | 0.003 |
| **PCR plate** | **3** | **1.501** | **0.029** | **0.014** |
| ***Individual ID*** | ***74*** | ***1.533*** | ***0.001*** | ***0.342*** |
| **Month** | **12** | **1.813** | **0.001** | **0.066** |
| Residual | 150 |  |  | 0.452 |
| Total | 240 |  |  | 1.000 |

**Table S4: The extent of individual-level variation (repeatability) in gut microbiota after controlling for known individual-level covariates.** Results are from a sequential PERMANOVA including all samples. A sequential PERMANOVA was used, in order to test the effect of individual identity once other (temporal and host) effects were accounted for. Terms are listed in the order they appear in the model, and significant terms (p<0.05) are shown in bold.

| **JACCARD** | | | | |
| --- | --- | --- | --- | --- |
|  | **Df** | **F** | **p** | **R^2^** |
| **Month** | **12** | **3.254** | **0.001** | **0.134** |
| **Age** | **2** | **1.492** | **0.013** | **0.010** |
| Sex | 1 | 1.390 | 0.067 | 0.005 |
| **Plot Region** | **8** | **1.705** | **0.001** | **0.047** |
| **Habitat type** | **3** | **1.713** | **0.001** | **0.018** |
| ***Individual ID*** | ***62*** | ***1.247*** | ***0.001*** | ***0.265*** |
| Residual | 152 |  |  | 0.521 |
| Total | 240 |  |  | 1.000 |
| **BRAY-CURTIS** | | | | |
|  | **Df** | **F** | **p** | **R^2^** |
| **Month** | **12** | **5.076** | **0.001** | **0.185** |
| **Age** | **2** | **1.821** | **0.011** | **0.011** |
| Sex | 1 | 1.639 | 0.053 | 0.005 |
| **Plot Region** | **8** | **2.118** | **0.001** | **0.051** |
| **Habitat type** | **3** | **2.295** | **0.001** | **0.021** |
| ***Individual ID*** | ***62*** | ***1.405*** | ***0.001*** | ***0.265*** |
| Residual | 152 |  |  | 0.462 |
| Total | 240 |  |  | 1.000 |

**Table S5: The effect of demographic individual-level variables on gut microbiota composition.** Results are from a marginal PERMANOVA on a dataset including one randomly selected sample per individual, with either Jaccard distance or Bray-Curtis dissimilarity as the response. Only one sample per ID was included so that the variance explained by individual-level factors could be directly compared (in a marginal PERMANOVA) without pseudoreplication. Significant terms (p<0.05) are shown in bold.

| **JACCARD INDEX** | | | | |
| --- | --- | --- | --- | --- |
|  | **Df** | **F** | **p** | **R^2^** |
| **Month** | **11** | **1.511** | **0.001** | **0.200** |
| Age | 2 | 1.222 | 0.090 | 0.029 |
| Sex | 1 | 1.037 | 0.332 | 0.012 |
| Region | 8 | 1.103 | 0.125 | 0.104 |
| **Habitat type** | **3** | **1.227** | **0.045** | **0.044** |
| Residual | 49 |  |  | 0.576 |
| Total | 74 |  |  | 1.000 |
| **BRAY-CURTIS** | | | | |
|  | **Df** | **F** | **p** | **R^2^** |
| **Month** | **11** | **1.858** | **0.001** | **0.221** |
| Age | 2 | 1.380 | 0.072 | 0.030 |
| Sex | 1 | 1.037 | 0.372 | 0.011 |
| Region | 8 | 1.132 | 0.151 | 0.098 |
| **Habitat type** | **3** | **1.371** | **0.050** | **0.045** |
| Residual | 49 |  |  | 0.531 |
| Total | 74 |  |  | 1.000 |

**Table S6: Correlations among social networks with varying edge definitions**. Results are shown from Mantel tests assessing the correlation between social networks varying in the edge definition used.

| **Network edge definition**  **(time window)** | **Correlation (r) with 12h network** | **p-value** |
| --- | --- | --- |
| 12h | 1 | 0.000 |
| 4h | 0.96 | 0.001 |
| 1h | 0.82 | 0.001 |
| 30min | 0.59 | 0.001 |
| 10min | 0.43 | 0.001 |
| 2min | 0.16 | 0.003 |

**Table S7: Variables predicting social association strength (Adjusted SRI) across networks.** Results of MRQAP models predicting Adjusted SRI for each social network. Significant terms (p<0.05) are shown in bold. Spatial distance significantly predicted social association strength in all networks, whereas kinship, age and sex similarity predicted social association strength in some, but not all, networks. More related mice were observed more often together in the 30min and 10min social network, adult mice were observed more often together than with juveniles in the 12h network, and opposite sex mice were more likely to be observed together in 30min network.

| **12h Social Network** | | | | | | |
| --- | --- | --- | --- | --- | --- | --- |
|  | **Estimate** | **P(β>=r)** | | | **P(β<=r)** | **P(\|β\|<=\|r\|)** |
| Intercept | 0.064 | 1.000 | | | 0.000 | 0.000 |
| Sex-similarity (0/1) | -0.003 | 0.115 | | | 0.885 | 0.267 |
| **Age-similarity (0/1)** | **0.009** | **0.989** | | | **0.011** | **0.041** |
| **Spatial distance** | **-0.121** | **0.000** | | | **1.000** | **0.000** |
| Kinship | 0.038 | 0.932 | | | 0.068 | 0.075 |
| **4h Social Network** | | | | | | |
|  | **Estimate** | | **P(β>=r)** | | **P(β<=r)** | **P(\|β\|<=\|r\|)** |
| intercept | 0.056 | | 1.000 | | 0.000 | 0.000 |
| Sex-similarity (0/1) | -0.004 | | 0.098 | | 0.902 | 0.204 |
| **Age-similarity (0/1)** | 0.006 | | 0.914 | | 0.086 | 0.174 |
| **Spatial distance** | **-0.101** | | **0.000** | | **1.000** | **0.000** |
| Kinship | 0.0372 | | 0.945 | | 0.055 | 0.058 |
| **1h Social Network** | | | | | | |
|  | **Estimate** | | **P(β>=r)** | | **P(β<=r** | **P(\|β\|<=\|r\|)** |
| intercept | 0.037 | | 1.000 | | 0.000 | 0.000 |
| Sex-similarity (0/1) | -0.004 | | 0.049 | | 0.951 | 0.094 |
| Age-similarity (0/1) | 0.001 | | 0.579 | | 0.421 | 0.793 |
| **Spatial distance** | **-0.063** | | **0.000** | | **1.000** | **0.000** |
| Kinship | 0.033 | | 0.938 | | 0.062 | 0.062 |
| **30min Social Network** | | | | | | |
|  | **Estimate** | | **P(β>=r)** | | **P(β<=r)** | **P(\|β\|<=\|r\|)** |
| intercept | 0.018 | | 1.000 | | 0.000 | 0.000 |
| **Sex-similarity (0/1)** | **-0.003** | | **0.005** | | **0.995** | **0.010** |
| Age-similarity (0/1) | 0.003 | | 0.972 | | 0.028 | 0.098 |
| **Spatial distance** | **-0.033** | | **0.000** | | **1.000** | **0.000** |
| **Kinship** | **0.039** | | **0.988** | | **0.012** | **0.012** |
| **10min Social Network** | | | | | | |
|  | **Estimate** | | **P(β>=r)** | | **P(β<=r)** | **P(\|β\|<=\|r\|)** |
| intercept | 0.008 | | 1.000 | | 0.000 | 0.000 |
| Sex-similarity (0/1) | -0.001 | | 0.288 | | 0.712 | 0.567 |
| Age-similarity (0/1) | -0.000 | | 0.276 | | 0.724 | 0.741 |
| **Spatial distance** | **-0.013** | | **0.000** | | **1.000** | **0.000** |
| **Kinship** | **0.036** | | **0.984** | | **0.016** | **0.016** |
| **2min Social Network** | | | | | | |
|  | **Estimate** | | | **P(β>=r)** | **P(β<=r)** | **P(\|β\|<=\|r\|)** |
| intercept | 0.001 | | | 0.986 | 0.014 | 0.014 |
| Sex-similarity (0/1) | -0.000 | | | 0.059 | 0.941 | 0.111 |
| Age-similarity (0/1) | 0.000 | | | 0.965 | 0.035 | 0.197 |
| **Spatial distance** | **-0.003** | | | **0.000** | **1.000** | **0.000** |
| Kinship | -0.001 | | | 0.298 | 0.702 | 0.412 |

**Table S8:** **Results of *brms* models testing the effect of social association strength and covariates on microbiota similarity (Jaccard Index).** Significant terms (where 95% credible intervals do not include zero) are shown in bold. Est.Error indicates the standard deviation of the posterior distribution.

| **12h Social Network** | | | | |
| --- | --- | --- | --- | --- |
|  | **Estimate** | **Est.Error** | **l-95% CI** | **u-95% CI** |
| Intercept | -1.46 | 0.06 | -1.57 | -1.35 |
| Sex | -0.00 | 0.01 | -0.02 | 0.01 |
| **Age** | **0.02** | **0.01** | **0.01** | **0.03** |
| Kinship | 0.00 | 0.04 | -0.08 | 0.09 |
| **Spatial distance** | **-0.08** | **0.02** | **-0.12** | **-0.04** |
| **Sampling interval** | **-0.46** | **0.01** | **-0.48** | **-0.43** |
| **Social Network** | **0.78** | **0.23** | **0.31** | **1.24** |
| **4h Social Network** | | | | |
|  | **Estimate** | **Est.Error** | **l-95% CI** | **u-95% CI** |
| Intercept | -1.45 | 0.05 | -1.55 | -1.34 |
| Sex | -0.00 | 0.01 | -0.02 | 0.01 |
| **Age** | **0.02** | **0.01** | **0.01** | **0.03** |
| Kinship | -0.01 | 0.04 | -0.09 | 0.08 |
| **Spatial distance** | **-0.09** | **0.02** | **-0.13** | **-0.04** |
| **Sampling interval** | **-0.47** | **0.01** | **-0.49** | **-0.44** |
| **Social Network** | **0.90** | **0.26** | **0.40** | **1.40** |
| **1h Social Network** | | | | |
|  | **Estimate** | **Est.Error** | **l-95% CI** | **u-95% CI** |
| Intercept | -1.44 | 0.05 | -1.55 | -1.33 |
| Sex | -0.00 | 0.01 | -0.02 | 0.01 |
| **Age** | **0.02** | **0.01** | **0.01** | **0.04** |
| Kinship | 0.01 | 0.04 | -0.07 | 0.10 |
| **Spatial distance** | **-0.09** | **0.02** | **-0.13** | **-0.05** |
| **Sampling interval** | **-0.48** | **0.01** | **-0.50** | **-0.46** |
| **Social Network** | **1.17** | **0.35** | **0.47** | **1.86** |
| **30min Social Network** | | | | |
|  | **Estimate** | **Est.Error** | **l-95% CI** | **u-95% CI** |
| Intercept | -1.43 | 0.05 | -1.54 | -1.33 |
| Sex | -0.00 | 0.01 | -0.02 | 0.01 |
| **Age** | **0.02** | **0.01** | **0.01** | **0.04** |
| Kinship | 0.01 | 0.04 | -0.08 | 0.09 |
| **Spatial distance** | **-0.09** | **0.02** | **-0.13** | **-0.05** |
| **Sampling interval** | **-0.49** | **0.01** | **-0.51** | **-0.46** |
| **Social Network** | **1.66** | **0.51** | **0.67** | **2.68** |
| **10min Social Network** | | | | |
|  | **Estimate** | **Est.Error** | **l-95% CI** | **u-95% CI** |
| Intercept | -1.41 | 0.05 | -1.52 | -1.30 |
| Sex | -0.00 | 0.01 | -0.02 | 0.01 |
| **Age** | **0.02** | **0.01** | **0.01** | **0.04** |
| Kinship | 0.01 | 0.05 | -0.08 | 0.10 |
| **Spatial distance** | **-0.12** | **0.02** | **-0.15** | **-0.08** |
| **Sampling interval** | **-0.50** | **0.01** | **-0.52** | **-0.48** |
| **Social Network** | **2.31** | **0.85** | **0.67** | **3.97** |
| **2min Social Network** | | | | |
|  | **Estimate** | **Est.Error** | **l-95% CI** | **u-95% CI** |
| Intercept | -1.40 | 0.05 | -1.51 | -1.29 |
| Sex | -0.00 | 0.01 | -0.02 | 0.01 |
| **Age** | **0.02** | **0.01** | **0.01** | **0.04** |
| Kinship | -0.00 | 0.04 | -0.09 | 0.09 |
| **Spatial distance** | **-0.15** | **0.02** | **-0.18** | **-0.11** |
| **Sampling interval** | **-0.51** | **0.01** | **-0.53** | **-0.49** |
| **Social Network** | **6.71** | **2.91** | **1.48** | **12.87** |

**Table S9**: **Results of MCMCglmm models testing the effect of social association strength and covariates on microbiota similarity.** Significant terms (p.MCMC<0.05) are shown in bold. post.mean indicates the mean of the posterior distribution of effect estimates, while l-95% CI and u-95% CI show the lower and upper limits of 95% credible intervals. Effective sample size (eff.samp) is the number of samples taken, adjusted for autocorrelation in the chain. An effect is considered significant if 95% credible intervals do not overlap zero, which is also indicated by p.MCMC-values. Specifically, p.MCMC values indicate the proportion of posterior samples (out of 1000) that fall on the other side of zero from the majority. The effect is considered significant when p.MCMC<0.05.

| **Jaccard Index** | | | | | | |
| --- | --- | --- | --- | --- | --- | --- |
|  | **post.mean** | | **l-95% CI** | **u-95% CI** | **eff.samp** | **p.MCMC** |
| (Intercept) | 0.206 | | 0.1938 | 0.2199 | 246 | <0.001 |
| Sex | -0.0004 | | -0.0023 | 0.0016 | 806 | 0.728 |
| **Age** | **0.0032** | | **0.0013** | **0.0052** | **874** | **0.002** |
| Kinship | 0.0015 | | -0.0107 | 0.0132 | 1163 | 0.830 |
| **Spatial distance** | **-0.0001** | | **-0.0001** | **-0.0000** | **1000** | **<0.001** |
| **Sampling interval** | **-0.0002** | | **-0.0002** | **-0.0002** | **1000** | **<0.001** |
| **12h Social Network** | **0.0629** | | **0.0505** | **0.0755** | **1000** | **<0.001** |
| **Bray-Curtis dissimilarity** | | | | | | |
|  | | **post.mean** | **l-95% CI** | **u-95% CI** | **eff.samp** | **p.MCMC** |
| (Intercept) | | 0.6692 | 0.6519 | 0.6898 | 647 | <0.001 |
| Sex | | 0.0007 | -0.0020 | 0.0032 | 1000 | 0.620 |
| **Age** | | **-0.0047** | **-0.0074** | **-0.0020** | **1200** | **<0.001** |
| Kinship | | -0.0009 | -0.018 | 0.016 | 1000 | 0.916 |
| **Spatial distance** | | **0.0001** | **0.0001** | **0.0002** | **1000** | **<0.001** |
| **Sampling interval** | | **0.0003** | **0.0003** | **0.0003** | **1286** | **<0.001** |
| **12h Social Network** | | **-0.0866** | **-0.1044** | **-0.0674** | **1000** | **<0.001** |

**T****able S10**: **Results of MRQAP models testing the effect of social association strength and covariates on microbiota similarity (Jaccard Index)**. Significant terms (mean p<0.05 across 100 iterations, each including a single randomly selected sample per individual) are shown in bold. Results are show both for the subset of individuals with kinship data, as well as the full dataset.

| Model term | Individuals with kinship data (n=70) | | All individuals  (n=75) | |
| --- | --- | --- | --- | --- |
|  | Mean Estimate | Mean  p-value | Mean Estimate | Mean  p-value |
| Sex-similarity (0/1) | -0.0022 | 0.53 | -0.0022 | 0.50 |
| Age-similarity (0/1) | -0.0004 | 0.50 | -0.0008 | 0.44 |
| Kinship | -0.0216 | 0.40 |  |  |
| Spatial distance | -0.0000 | 0.41 | -0.0000 | 0.41 |
| **Sampling interval** | **-0.0002** | **0.00** | **-0.0002** | **0.00** |
| 12h Social Network | 0.0624 | 0.06 | **0.07** | **0.02** |

**Table S11:** **Results of *brms* models testing the effect of binary social association (BI) and covariates on microbiota similarity (Jaccard Index).** Significant terms (where 95% credible intervals do not include zero) are shown in bold.

| **12h Social Network** | | | | |
| --- | --- | --- | --- | --- |
|  | **Estimate** | **Est.Error** | **l-95% CI** | **u-95% CI** |
| Intercept | -1.47 | 0.06 | -1.58 | -1.36 |
| Age | 0.01 | 0.01 | 0.00 | 0.03 |
| Kinship | 0.01 | 0.04 | -0.08 | 0.10 |
| Sex | -0.00 | 0.01 | -0.01 | 0.01 |
| **Spatial distance** | **-0.10** | **0.02** | **-0.13** | **-0.06** |
| **Sampling interval** | **-0.43** | **0.01** | **-0.45** | **-0.40** |
| **12h Social Network** | **0.16** | **0.04** | **0.08** | **0.24** |
| **4h Social Network** | | | | |
|  | **Estimate** | **Est.Error** | **l-95% CI** | **u-95% CI** |
| Intercept | -1.46 | 0.06 | -1.58 | -1.35 |
| Age | 0.01 | 0.01 | 0.00 | 0.03 |
| Kinship | -0.00 | 0.04 | -0.09 | 0.08 |
| Sex | -0.00 | 0.01 | -0.02 | 0.01 |
| **Spatial distance** | **-0.10** | **0.02** | **-0.13** | **-0.06** |
| **Sampling interval** | **-0.44** | **0.01** | **-0.47** | **-0.41** |
| **4h Social Network** | **0.15** | **0.04** | **0.07** | **0.23** |
| **1h Social Network** | | | | |
|  | **Estimate** | **Est.Error** | **l-95% CI** | **u-95% CI** |
| Intercept | -1.45 | 0.06 | -1.56 | -1.34 |
| Age | 0.02 | 0.01 | 0.00 | 0.03 |
| Kinship | -0.01 | 0.05 | -0.10 | 0.08 |
| Sex | -0.00 | 0.01 | -0.02 | 0.01 |
| **Spatial distance** | **-0.09** | **0.02** | **-0.13** | **-0.05** |
| **Sampling interval** | **-0.47** | **0.01** | **-0.49** | **-0.44** |
| **1h Social Network** | **0.14** | **0.04** | **0.06** | **0.21** |
| **30min Social Network** | | | | |
|  | **Estimate** | **Est.Error** | **l-95% CI** | **u-95% CI** |
| Intercept | -1.44 | 0.05 | -1.55 | -1.34 |
| **Age** | **0.02** | **0.01** | **0.01** | **0.03** |
| Kinship | 0.00 | 0.05 | -0.09 | 0.09 |
| Sex | -0.00 | 0.01 | -0.02 | 0.01 |
| **Spatial distance** | **-0.09** | **0.02** | **-0.12** | **-0.05** |
| **Sampling interval** | **-0.48** | **0.01** | **-0.50** | **-0.45** |
| **30min Social Network** | **0.14** | **0.04** | **0.06** | **0.21** |
| **10min Social Network** | | | | |
|  | **Estimate** | **Est.Error** | **l-95% CI** | **u-95% CI** |
| Intercept | -1.42 | 0.05 | -1.52 | -1.32 |
| **Age** | **0.02** | **0.01** | **0.01** | **0.04** |
| Kinship | 0.03 | 0.04 | -0.06 | 0.11 |
| Sex | -0.00 | 0.01 | -0.01 | 0.01 |
| **Spatial distance** | **-0.11** | **0.02** | **-0.15** | **-0.07** |
| **Sampling interval** | **-0.50** | **0.01** | **-0.52** | **-0.48** |
| **10min Social Network** | **0.13** | **0.04** | **0.06** | **0.20** |
| **2min Social Network** | | | | |
|  | **Estimate** | **Est.Error** | **l-95% CI** | **u-95% CI** |
| Intercept | -1.40 | 0.05 | -1.51 | -1.30 |
| **Age** | **0.03** | **0.01** | **0.01** | **0.04** |
| Kinship | 0.00 | 0.04 | -0.09 | 0.09 |
| Sex | -0.00 | 0.01 | -0.01 | 0.01 |
| **Spatial distance** | **-0.13** | **0.02** | **-0.17** | **-0.10** |
| **Sampling interval** | **-0.51** | **0.01** | **-0.53** | **-0.49** |
| **2min Social Network** | **0.14** | **0.05** | **0.05** | **0.25** |

**T****able S12:** **Results of *brms* models testing whether the effect of social association strength on microbiota similarity (Jaccard Index) varies according to the sex of interacting individuals.**

| **12h Social Network** | | | | |
| --- | --- | --- | --- | --- |
|  | **Estimate** | **Est.Error** | **l-95% CI** | **u-95% CI** |
| Intercept | -1.51 | 0.09 | -1.69 | -1.34 |
| **Age** | **0.02** | **0.01** | **0.01** | **0.04** |
| Kinship | -0.00 | 0.04 | -0.09 | 0.09 |
| Spatial distance | -0.09 | 0.02 | -0.13 | -0.05 |
| Sampling interval | -0.49 | 0.01 | -0.52 | -0.47 |
| Sex category: male-male | 0.12 | 0.11 | -0.10 | 0.33 |
| Sex category: female-male | 0.06 | 0.06 | -0.05 | 0.17 |
| Social Network | 0.10 | 0.13 | -0.15 | 0.35 |
| Social Network: Sex category, male-male | 0.28 | 0.14 | 0.01 | 0.56 |
| Social Network: Sex category, male-female | 0.30 | 0.13 | 0.04 | 0.56 |

**Table S13: Results of MCMCglmm models predicting microbiota diversity with covariates**. Results from simplified MCMCglmm models predicting either asymptotic microbial Shannon diversity or asymptotic richness as a function of temporal, individual host and methodological factors. Models were simplified to remove the non-significant (p>0.05) variables age and sex. Significant (p<0.05) terms are shown in bold.

| **Asymptotic Shannon Diversity** | | | | | |
| --- | --- | --- | --- | --- | --- |
|  | **post.mean** | **l-95% CI** | **u-95% CI** | **eff.samp** | **p.MCMC** |
| (Intercept) | 83.55 | 48.71000 | 117.94433 | 1000.0 | <0.001 |
| month2 | 15.19 | -2.16278 | 34.97057 | 1269.9 | 0.108 |
| month3 | 10.98 | -8.55660 | 25.70075 | 1000.0 | 0.188 |
| **month4** | **18.48** | **0.75664** | **34.82477** | **888.5** | **0.024** |
| month5 | 15.51 | -5.66490 | 35.37272 | 1000.0 | 0.162 |
| month6 | 2.08 | -14.00984 | 17.46371 | 911.8 | 0.810 |
| month7 | 2.37 | -24.16373 | 33.97004 | 1000.0 | 0.856 |
| **month8** | **24.96** | **8.70720** | **43.91518** | **902.6** | **0.006** |
| month9 | 7.01 | -14.94552 | 27.57888 | 809.6 | 0.514 |
| month10 | 13.90 | -1.87155 | 29.43618 | 1169.5 | 0.102 |
| month11 | 0.88 | -10.47376 | 12.65884 | 1000.0 | 0.900 |
| month12 | 10.81 | -7.44916 | 27.69678 | 1000.0 | 0.232 |
| **pcr_plate2** | **18.37** | **9.29907** | **27.59661** | **1000.0** | **<0.001** |
| pcr_plate3 | 6.13 | -2.55688 | 15.49759 | 1000.0 | 0.186 |
| pcr_plate5 | 5.06 | -12.85787 | 23.24887 | 1000.0 | 0.596 |
| **Habitat_Mixed** | **-58.88** | **-92.06864** | **-25.08360** | **1144.8** | **0.002** |
| **Habitat_Open_woodland** | **-57.60** | **-88.02739** | **-26.77238** | **1146.2** | **0.002** |
| **Habitat_Rhododendron** | **-51.57** | **-82.94973** | **-18.69522** | **1184.4** | **0.002** |
| Region2 | -7.88 | -21.26818 | 6.96602 | 1000.0 | 0.304 |
| Region3 | 14.64 | -2.69574 | 32.01329 | 1000.0 | 0.090 |
| Region4 | -6.85 | -24.49686 | 8.85641 | 927.1 | 0.444 |
| Region5 | 1.75 | -15.79396 | 19.06128 | 460.8 | 0.874 |
| Region6 | -0.07 | -17.90074 | 15.07547 | 797.1 | 0.976 |
| Region7 | 12.69 | -4.78729 | 29.14240 | 886.3 | 0.136 |
| Region8 | -1.42 | -26.49774 | 25.18387 | 1132.5 | 0.906 |
| Region9 | 3.05 | -10.21841 | 19.17771 | 1000.0 | 0.698 |
| **Asymptotic richness** | | | | | |
|  | **post.mean** | **l-95% CI** | **u-95% CI** | **eff.samp** | **p.MCMC** |
| (Intercept) | 0.018 | 1.089e+02 | 2.713e+02 | 1000 | <0.001 |
| **Read Count** | **0.003** | **1.676e-03** | **4.016e-03** | **1000** | **<0.001** |
| **month2** | **0.466** | **8.656e+00** | **8.777e+01** | **1000** | **0.030** |
| **month3** | **0.398** | **7.099e+00** | **7.081e+01** | **1000** | **0.016** |
| **month4** | **0.419** | **7.615e+00** | **8.403e+01** | **1173** | **0.030** |
| month5 | 0.237 | -2.304e+01 | 6.739e+01 | 1000 | 0.308 |
| month6 | -0.175 | -5.028e+01 | 1.863e+01 | 1000 | 0.342 |
| month7 | 0.130 | -4.848e+01 | 7.605e+01 | 1000 | 0.712 |
| month8 | 0.297 | -1.185e+01 | 6.396e+01 | 1000 | 0.148 |
| month9 | -0.246 | -7.079e+01 | 1.649e+01 | 1124 | 0.268 |
| month10 | -0.142 | -4.616e+01 | 1.831e+01 | 1000 | 0.392 |
| month11 | 1.465 | -2.417e+01 | 2.552e+01 | 1000 | 0.896 |
| **month12** | **0.378** | **8.248e-01** | **7.357e+01** | **1000** | **0.046** |
| **pcr_plate2** | **0.324** | **1.141e+01** | **5.155e+01** | **1222** | **0.006** |
| pcr_plate3 | 6.562 | -1.347e+01 | 2.638e+01 | 1000 | 0.518 |
| pcr_plate5 | -0.207 | -6.065e+01 | 1.664e+01 | 1000 | 0.292 |
| Habitat_mixed | -0.599 | -1.387e+02 | 2.744e+00 | 1075 | 0.084 |
| Habitat_open_woodland | -0.558 | -1.171e+02 | 2.261e+00 | 1000 | 0.076 |
| Habitat_Rhododendron | -0.514 | -1.137e+02 | 1.160e+01 | 1000 | 0.130 |
| Region2 | -0.971 | -3.852e+01 | 2.235e+01 | 1000 | 0.526 |
| **Region3** | **0.387** | **3.255e+00** | **7.444e+01** | **1000** | **0.036** |
| Region4 | -0.584 | -3.967e+01 | 2.973e+01 | 1000 | 0.724 |
| Region5 | -0.134 | -4.988e+01 | 2.543e+01 | 1000 | 0.508 |
| Region6 | -1.631 | -3.515e+01 | 3.077e+01 | 1000 | 0.948 |
| Region7 | 7.049 | -2.751e+01 | 3.996e+01 | 1000 | 0.674 |
| Region8 | 7.802 | -4.797e+01 | 5.899e+01 | 1000 | 0.750 |
| Region9 | 0.142 | -1.771e+01 | 4.365e+01 | 1000 | 0.374 |

**Table S14: Effects of social centrality metrics and non-social factors on gut microbiota Shannon diversity**

Summary of social centrality metric effects on microbiota Shannon diversity, in MCMCglmm models including a single social centrality measure and controlling for all significant covariates identified in Table S13. Significance is inferred with p.MCMC, a Bayesian p-value describing the extent to which credible intervals of posterior samples overlap zero.

|  | **12h network** | | **2min network** | |
| --- | --- | --- | --- | --- |
|  | **Posterior mean**  **(95% credible interval)** | **p.MCMC** | **Posterior mean**  **(95% credible interval)** | **p.MCMC** |
| **Intercept** | 6.4 |  | 4.7 |  |
| **Degree** | 0.017 (CI -0.011-0.047) | 0.272 | 0.102 (CI -0.032-0.227) | 0.150 |
| **Weighted degree** | -0.014 (CI -0.165-0.115) | 0.864 | 0.032 (CI -0.127-0.172) | 0.670 |
| **EVC** | 0.017 (CI -0.135-0.159 | 0.830 | 0.004 (CI -0.162-0.162) | 0.999 |
| **Betweenness** | 0.040 (CI -0.071-0.138) | 0.472 | 0.074 (CI -0.042-0.175) | 0.188 |
| **Information Centrality** | 0.070 (CI- 0.062-0.180) | 0.266 | 0.090 (CI -0.021-0.224) | 0.144 |
| **Bridge Propensity** | 0.073 (CI -0.160-0.309) | 0.546 | 0.060 (CI -0.037-0.158) | 0.240 |

**Supplementary References**

1.	Godsall B, Coulson T, Malo AF. From physiology to space use: energy reserves and androgenization explain home-range size variation in a woodland rodent. Montgomery I, editor. *J Anim Ecol* 2014; 83(1):126–35.

2.	Wilson A. Landscape genetics of highly disturbed arable systems: Insights gained from investigating a small mammal species*.* PhD thesis. The University of St. Andrews. 2014

3.	Godsall B. Mechanisms of space use in the wood mouse, Apodemus sylvaticus. PhD thesis. Imperial College London. 2015

4.	Kalinowski ST, Taper ML, Marshall TC. Revising how the computer program CERVUS accommodates genotyping error increases success in paternity assignment. Molecular ecology. 2007 Mar;16(5):1099-106.

5.	Van Oosterhout C, Hutchinson WF, Wills DP, Shipley P. MICRO‐CHECKER: software for identifying and correcting genotyping errors in microsatellite data. *Molecular Ecology Notes* 2004 Sep;4(3):535-8.

6.	Wang J, Santure AW. Parentage and sibship inference from multilocus genotype data under polygamy. *Genetics* 2009 Apr 1;181(4):1579-94.

7.	Jones OR, Wang J. COLONY: a program for parentage and sibship inference from multilocus genotype data. *Mol Ecol Resour* 2010; 10(3):551–5.

8.	Caporaso JG, Lauber CL, Walters WA, Berg-Lyons D, Lozupone CA, Turnbaugh PJ, *et al.* Global patterns of 16S rRNA diversity at a depth of millions of sequences per sample. *Proc Natl Acad Sci USA* 2011; 108:4516–22.

9.	D’Amore R, Ijaz UZ, Schirmer M, Kenny JG, Gregory R, Darby AC, *et al.* A comprehensive benchmarking study of protocols and sequencing platforms for 16S rRNA community profiling. *BMC Genomics* 2016; 17(1):55.

10.	<https://support.illumina.com/content/dam/illumina-support/documents/documentation/chemistry_documentation/samplepreps_nextera/nexteradna/nextera-dna-library-prep-reference-guide-15027987-01.pdf>

11.	Callahan BJ, McMurdie PJ, Rosen MJ, Han AW, Johnson AJ, Holmes SP. DADA2: high-resolution sample inference from Illumina amplicon data. *Nature methods.* 2016 Jul;13(7):581-3.

12.	Martin M. Cutadapt removes adapter sequences from high-throughput sequencing reads. *EMBnet. journal* 2011 May 2;17(1):10-2.

13.	McMurdie PJ, Holmes S. phyloseq: An R Package for Reproducible Interactive Analysis and Graphics of Microbiome Census Data. *PLoS One* 2013; 8(4)

14.	Hsieh TC, Ma KH, Chao A. iNEXT: an R package for rarefaction and extrapolation of species diversity ( Hill numbers). *Methods Ecol Evol* 2016; 7(12):1451–6.

15.	McKnight DT, Huerlimann R, Bower DS, Schwarzkopf L, Alford RA, Zenger KR. Methods for normalizing microbiome data: An ecological perspective. *Methods Ecol Evol* 2019; 10(3):389–400

16.	Hadfield JD. MCMC methods for multi-response generalized linear mixed models: the MCMCglmm R package. *Journal of statistical software* 2010 Feb 2;33(2):1-22.

17.	Dekker D, Krackhardt D, Snijders TA. Sensitivity of MRQAP tests to collinearity and autocorrelation conditions. *Psychometrika* 2007 Dec 1;72(4):563-81

18.	Shizuka D, Chaine AS, Anderson J, Johnson O, Laursen IM, Lyon BE. Across-year social stability shapes network structure in wintering migrant sparrows. *Ecol Lett* 2014;17(8):998–1007

19.	Firth JA, Sheldon BC. Social carry‐over effects underpin trans‐seasonally linked structure in a wild bird population. *Ecology letters* 2016 Nov;19(11):1324-32.
